# Resolving cell lineages and gene functions in the developing mouse gastrointestinal tract using *in utero* transduction

**DOI:** 10.64898/2026.04.09.716777

**Authors:** Ziwei Liu, Krishnanand Padmanabhan, Jingyan He, Katrin Hector, Bettina Semsch, Jia Sun, Viktoria Knoflach, Sarantis Giatrellis, Khachatur Dallakyan, Christian Göritz, Emma Rachel Andersson, Ulrika Marklund

## Abstract

How diverse cell lineages emerge and are genetically regulated during organogenesis are central questions in understanding the developmental origins of disease. However, the mouse gut, including its intrinsic enteric nervous system (ENS) derived from migratory neural crest, has remained difficult to experimentally target. Here, we introduce an *in utero* lentiviral nano-injection strategy that enables early and efficient access to progenitor cells of all major cell types within the developing gut as well as gut-innervating ganglia. Leveraging this approach in combination with DNA barcoding and single cell transcriptomics, we resolve clonal relationships in all gut lineages, including epithelial, neural, immune, and mesenchymal cell types. Clonal coupling between distinct subsets of fibroblasts and either pericytes, mesothelial cells, or interstitial cells of Cajal, suggested a developmental logic whereby the mesenchymal compartment arises from a set of fate-biased progenitors. Yet, mesenchymal regionalization along the anterior–posterior axis establishes early, whereas the ENS displays broad clonal dispersion across gut regions and acquires subsequent regional identities. We further adapted the platform for temporally controlled cell-type specific gene manipulation and, as a proof-of-principle, show that induced expression of the proneural factor Ascl1 biases ENS progenitor cells toward neuronal differentiation. Together, this work provides insights into refined spatiotemporal lineage relationships within a multigerm-layer organ and establishes a broadly applicable *in vivo* framework for probing gene function during gastrointestinal and neural crest development.

**SIGNIFICANCE:** The gastrointestinal tract comprises diverse cell types originating from all three germ layers and includes the neural crest-derived enteric nervous system (ENS). Progress in defining these lineages and their gene regulation is challenged by the limited experimental access to the developing gut. Here, we establish *in utero* lentiviral transduction as an efficient approach to resolve clonal lineages and address gene functions in defined gut cell types. We show that mesenchyme assumes positional allocation early and differentiates through fate-restricted progenitors, linking specialized mesenchymal cell types to different fibroblasts. In contrast, the ENS differentiates stochastically and acquires late regional identities. Our study reveals fundamental principles of multi-lineage organogenesis and provides a framework to dissect the contribution of developmental programs to visceral dysfunction.

## INTRODUCTION

The gastrointestinal (GI) tract is essential for life. Congenital gut malformations including Hirschsprung disease, pyloric stenosis, and various intestinal atresias result in diverse and life-long complications. In the general population, 30% suffer from chronic gut dysfunctions, including disorders of gut-brain interaction (DGBIs) and inflammatory bowel disease (IBD) (1). Conceivably these conditions may arise from subtle alterations in gut cell composition already established at fetal stages (2). Supporting this notion, premature birth is an identified risk factor for developing DGBIs (3). Furthermore, congenital chronic intestinal pseudo-obstruction (CIPO) can be myopathic, mesenchymopathic or neuropathic (4), underscoring the critical role of coordinated interactions among multiple gut cell types in maintaining GI motility. Thus, there is a need to better resolve how different cell lineages in the GI-tract develop and integrate during early life.

Experimental gene perturbation/manipulation - genetic gain or loss of function – are essential tools to resolve how gene regulatory networks control cellular differentiation and how developmental deviations can lead to pathology. Owing to the relative ease with which the central nervous system (CNS) can be genetically manipulated using *in vivo* embryonic transduction or electroporation (5), the CNS field has gained intricate knowledge of precise gene regulatory networks controlling the generation of different neuronal subtypes (6, 7). These manipulations are possible due to the ventricular structure of the brain and the lumen of the spinal cord, making them accessible to plasmids or viral vectors. In contrast, the GI-tract is positioned internally and not possible to target using these common approaches.

Normal gut physiology is critically dependent on neuronal control provided by its intrinsic enteric nervous system (ENS), essential for peristalsis, secretion, and blood flow regulation. These functions are modulated by sympathetic innervation, primarily of celiac ganglion neurons, while sensory information is captured from the gut via dorsal root ganglia (DRG) at the trunk regions, and nodose ganglia in cranial regions. The ENS, sympathetic ganglia, and DRG are formed from neural crest cells (NCC), a transient embryonic population originating from the dorsal neural tube (8). In the mouse, delamination of NCC occurs around embryonic day (E)8 (Theiler stages TS12-13) and thereafter, the migrating cells and the ganglia that form are positioned unfavorably for traditional *in utero* manipulations that are typically performed from E11.5 (5). Thus, experimental systems to rapidly investigate the development of the murine gut, the ENS, and other gut-innervating ganglia have been lacking.

Recently, we developed neural plate targeting with *in utero* nano-injection (NEPTUNE) to target the developing CNS at earlier stages (9, 10). With NEPTUNE, ultrasound is used to guide lentiviral delivery of cargo into the mouse amniotic cavity as early as E7.5, allowing access to 80-95% of cells in the CNS. The technique can also be used to introduce high-diversity lentiviral barcode libraries to resolve refined lineages using next generation single cell lineage tracing (11, 12). As distinct barcodes integrate in the DNA of each transduced cell, clonal relationships can be reconstructed after successive cell divisions by single cell RNA-sequencing (scRNA-seq), revealing how individual progenitors contribute to the generation of diverse cell types. Here, we asked whether the method allows for targeting of the GI-tract and of neural crest/placodes, whose derivatives innervate the gut, for the purpose of revealing clonally linked gut cell types and performing genetic experimentation of the developing ENS.

We found that ultrasound-guided *in utero* lentiviral nano-injection at E7.5 transduces progenitors that contribute to the GI tract, including endodermal (i.e. epithelia), mesodermal (e.g. smooth muscle cells, fibroblasts and interstitial cells of Cajal; ICCs) and ectodermal progenitors (e.g. ENS). In addition, the method efficiently targets extrinsic gut-innervating ganglia (e.g DRG). By combining high-throughput clonal labeling and transcriptomic profiling of cells in the gut, we identified fate-restricted progenitor cells present at the time of transduction and reconstructed lineage relationships across differentiating gut cell types and regions. Of note, we uncovered molecularly defined fibroblast populations that are clonally coupled with either mesothelial cells, pericytes or ICCs, revealing early diversification paths within the gut mesenchyme. We find that ENS targeting is most efficient at early E7.5, but that residual epiblast cells are still present at this stage. Beyond lineage tracing, we show that *in utero* nano-injections can be utilized for conditional gene perturbation in the ENS. Using the pan-neuronal Baf53-Cre mouse line, we demonstrate that lentivirus carrying Cre-dependent cargo is selectively expressed in developing enteric neurons. In Sox10-CreERT2 mice, transgene expression can instead be induced in ENS progenitors/glia in a temporally controlled manner. As proof-of-principle, we show that timed overexpression of the proneural transcription factor *Ascl1* promote neuronal differentiation. Taken together, our results show that ultrasound-guided *in utero* nano-injection is a versatile tool to investigate organogenesis and ENS development. Our work offers insight into how neural crest and mesoderm lineages diversify and regionally pattern within the developing gut.

## MATERIAL and METHODS

### Mice

The following transgenic and reporter mouse strains were used: Baf53b-Cre (JAX, #027826), Sox10-CreERT2 (SER26) (13), R26-EYFP (JAX #006148) and R26-tdTomato (Ai14; JAX #007908). C57/B6J and CD1 (Charles River Laboratories) wild-type (WT) mice were used. Animals were group-housed, with food and water *ad libitum,* under 12-h light-dark cycle conditions, 22°C ambient temperature and 50% humidity. Animal experiments were approved by the local ethics committee (Stockholm Norra djurförsöksetiska nämnd, Jordbruksverket; N5264/18, N6626-2019, N5237-2023, 8188-2017, 2987-2020 and 315-23).

### Lentivirus constructs

Plasmid *LV-H2B-GFP* was a gift from Elaine Fuchs (Addgene #25999). *LV-EF1A-H2B-tdTomato-30N* was previously described (11, 12). *Lenti-DIO-EGFP* (Supplementary Figure 1A) was based on the interim vector *pRRLSIN-cPPT-hPGK-EGFP-WPRE* (Gift from Didier Trono, Addgene #12252). *Lenti-DIO-EGFP-Ascl1* (Supplementary Figure 1B) was constructed based on *pRRLSIN-cPPT-shortCAG-dlox-EGFP_2A(PTV)(rev)-dlox-WPRE* and the GeneArt product *mAscl1* (710bp). Both vectors were constructed, amplified, and characterized by the Viral Vector Facility (VVF) of the Neuroscience Center Zurich (ZNZ). Plasmids and virus were produced and characterized as described in (Supplementary Material and Methods). High-titer vesicular stomatitis virus G (VSV-G)–pseudotyped lentiviral particles were produced either in-house or by a commercial facility (GEG TECH, France) and used at a minimum titer of 1 × 10⁹ transducing units (TU)/mL.

### Ultrasound-guided *in utero* nano-injections

A detailed description of the procedure has been reported (9, 10) and is described in (Supplementary Material and Methods). We considered Theiler stages TS11A–C to represent the optimal E7.5 window for ultrasound-guided *in utero* injections (Supplementary Figure 4A, B), while injection at E8 or later failed to efficiently target gut and ENS (Supplementary Figure 4C, D).

### Tamoxifen administration

Plug-mated females received tamoxifen (0.1 mg/g body weight; Sigma) dissolved in corn oil/ethanol (9:1) by intraperitoneal injection at E16.0-E16.5. Embryos were collected 26 or 55 hours after tamoxifen administration.

### Tissue preparation and immunohistochemistry

E13.5 embryos were decapitated and fixed in 4% PFA in PBS for 4-5 hours at 4 °C. Intestines from E14.5-E18.5 were dissected out and fixed in 4% PFA in PBS at 4 °C for 1.5-2 hours. Samples were washed in PBS three times and dehydrated in 30% sucrose in PBS at 4 °C overnight. Intestines were cut into 1 cm long pieces, representing different regions and embedded for transverse analysis in OCT freezing medium and stored at -80 °C. Samples were sectioned at 12 μm and stored at -20 °C.

Frozen tissue sections were air-dried at RT for 30 min, rinsed with PBS and incubated in blocking solution containing purified donkey anti-mouse Fab (Jackson Laboratories, PA, USA #715-007-003) diluted 1:50 in PBS for 2 hours at room temperature (RT). After rinsing with PBS, the sections were blocked with 2% normal donkey serum (NDS, Jackson) and 0.1% Triton X-100 (Sigma) in PBS for 2 hours, and incubated with primary antibodies diluted in the same solution overnight at 4°C. They were then washed three times with PBS (10 minutes each) and incubated with secondary antibodies at RT for 1 hour. Tissue was washed three times with PBS (10 minutes each) and mounted in DAKO mounting medium (Agilent) containing DAPI (4’,6-diamidino-2-phenylindole). Primary and secondary antibodies are specified in (Supplementary Material and Methods.

### Preparation of single-cell RNA sequencing datasets

CD1 embryos injected with LV-EF1A-H2B-tdTomato-30N were harvested at E16.5. Stomach and 1/3 of the distal small intestine were harvested from the E7.5 experiment, while the stomach, 2/3 of the small intestine, and the entire large intestine were harvested from the E7.5^Early^ experiment. Dissociation was performed as described in (Supplementary Material and Methods). Briefly, tissues were cut, enzymatically processed, triturated and TOM+ DRAQ7-cells isolated through flow cytometry (Supplementary Figure 1C-H). cDNA libraries were prepared using 10x Chromium Next GEM Single Cell 3’ Reagent Kits Version 3.1 Dual Index (10x Genomics, CA, USA). The datasets were named according to stage of injection (i) and analysis (a): iE7.5-aE16.5 and iE7.5^Early^-aE16.5.

### Analysis of single-cell RNA sequencing datasets

A detailed description of the scRNA-seq analysis can be found in (Supplementary Material and Methods). In brief, E16.5 scRNA-seq datasets were aligned to a combined mouse genome (GRCm38) with a tdTomato-N transgene containing a 30bp barcode region (11, 12) using CellRanger. Raw count matrices were processed in Seurat, with ambient RNA contamination assessed and corrected using SoupX. Low-quality cells and doublets were removed based on gene/UMI thresholds, mitochondrial contents and DoubletFinder predictions. Data were normalized using SCTransform, followed by dimensionality reduction, batch correction with Harmony, clustering and UMAP visualisation. CloneIDs were assigned to cells using the TREX (TRacking and gene EXpression profiling of clonally related cells) pipeline (11, 12) based on shared barcodes. Clonal composition and relationships between cell types were analyzed using statistical comparisons to randomized data, generating clonal coupling scores and correlation-based clustering to infer lineage relationships.

### Data Accession

scRNA-seq data used for analysis in Figure 1 and Supplementary Figure 2 can be found at ArrayExpress with the accession number E-MTAB-14817 (11). The scRNA-seq data generated in this study will be made available at GEO upon publication.

**Figure 1.**
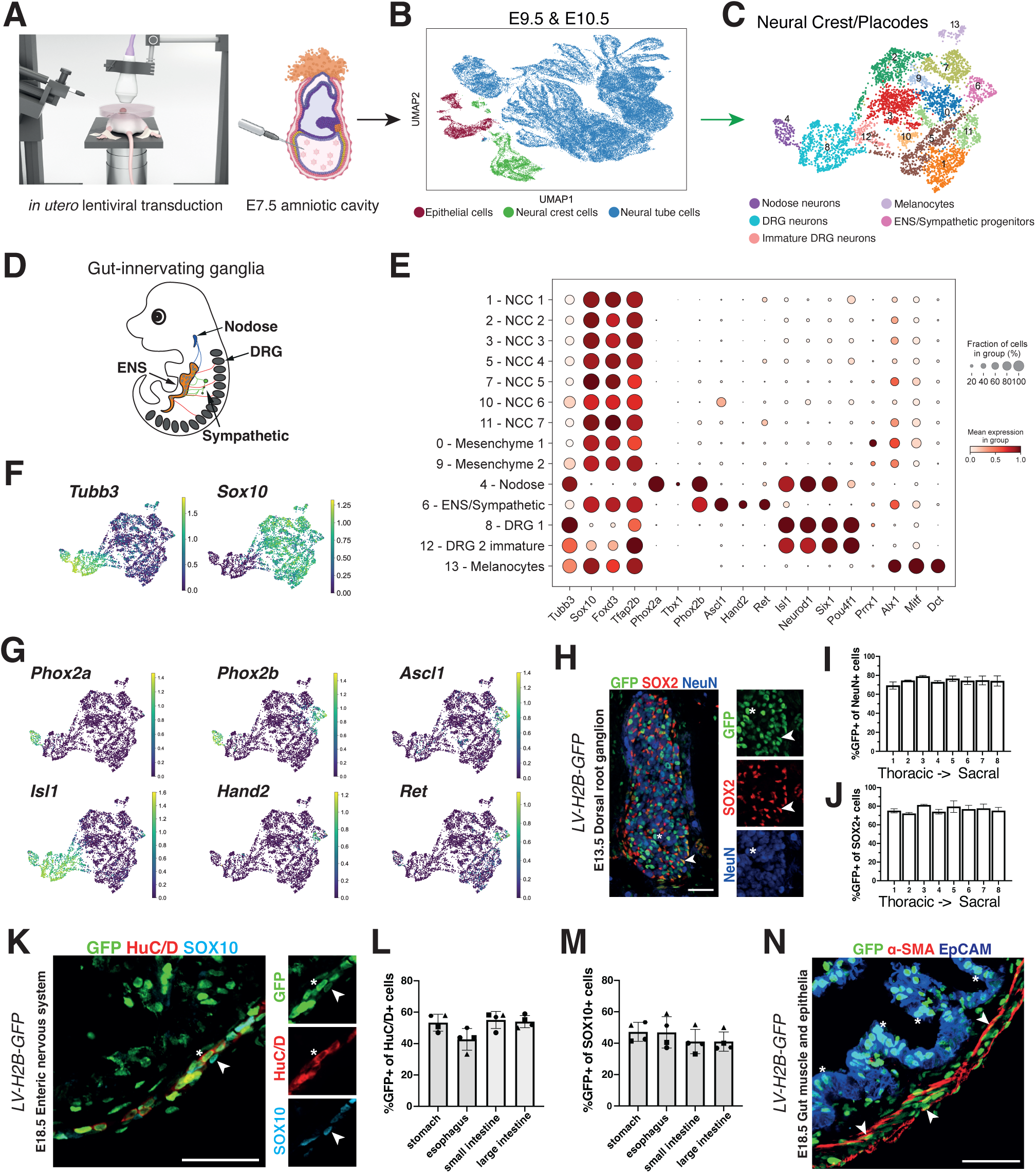
Efficient targeting of neural crest/placode populations destined to innervate the gut by *in utero* transduction at E7.5. (A) Schematic drawing of experimental setup for ultrasound-guided *in utero* nano-injection. Left-most panel reproduced from Mangold et al 2021 (10), with permission. The right panel was created with BioRender.com. (B) Uniform manifold approximation and projection (UMAP) plot of cells recovered from E9.5 and E10.5 embryos following E7.5 amniotic cavity injection (11). (C) UMAP plot of reprocessed neural crest/placode cells (green populations in B), revealing 14 clusters. (D) Schematic drawing of neural crest/placode-derived ganglia innervating the gut. (E) Dot plot showing expression of canonical marker genes for neural crest/placode populations across the clusters. Dot size represents the fraction of cells expressing the gene and color intensity represents mean expression. (F) Feature plots showing expression of neuronal (*Tubb3+*) and migratory neural crest/placode (*Sox10+*) markers. (G) Feature plots showing gene expression associated with nodose ganglia, sympathetic ganglia, DRG and the ENS. (J) Representative immunofluorescence staining of an E13.5 DRG showing H2B-GFP expression in both SOX2+ neural progenitors (arrowheads) and NeuN+ postmitotic neurons (asterisks). (I, J) Bar graphs showing proportions of GFP positive NeuN+ neurons and SOX2+ neural progenitors along the thoracic-sacral axis (n=3 mice). Bars represent the mean ± standard deviation (SD). (K) Representative immunofluorescence image of E18.5 small intestine showing GFP expression in HuC/D+ neurons (asterisks) and SOX10+ ENS progenitors and glia (arrowheads). (L, M) Bar graphs showing proportion of GFP positive HuC/D+ neurons and SOX10+ progenitors/glia across gut regions (n=4 mice). Bars represent mean ± SD. Each dot represents one animal. (N) Representative immunofluorescence image of E18.5 gut showing GFP expression in alpha-SMA+ smooth muscle (arrowheads) and EpCAM+ epithelium (asterisks). Scale bars: 50µm.

## RESULTS

### Efficient targeting of gut-innervating ganglia by *in utero* transduction at E7.5

Previous studies have demonstrated that high-titer lentivirus injection into the amniotic cavity of E7.5 embryos results in widespread transduction of the neural plate and otic placodes (9–11). Because gut-innervating ganglia originate from neural crest or from placodes, we hypothesized that these structures could also be targeted. To assess whether E7.5 amniotic cavity lentiviral injection targets neural plate progenitors that generate gut-innervating neuronal populations (Figure 1A), we re-analyzed a published scRNA-seq dataset of E9.5 and E10.5 embryos injected at E7.5 with lentiviruses carrying *tdTomato* (11). The cells in this dataset had been broadly categorized as “neural tube cells”, “epithelial cells”, and “neural crest cells” (Figure 1B, Supplementary Figure 2A). To investigate whether developing gut-innervating neurons had been targeted (Figure 1D), we renormalized and subclustered the “neural crest cells” (4,916 cells), which yielded 14 clusters (Figure 1C, E; Supplementary Figure 2B). We identified three *Tubb3+* neuronal clusters (clusters 4, 8, 12), while remaining clusters displayed high *Sox10* expression, indicative of neural progenitor or other immature cell states (Figure 1E, F; Supplementary Figure 2C, D). Amongst *Sox10*+ clusters, clusters 0 and 9 were defined as mesenchymal as they expressed *Prrx1* and *Alx1*, while cluster 13 showed a melanocyte identity with high *Mitf* and *Dct* expression. Cluster 6 exhibited an autonomic nervous system character (*Phox2b*+, *Hand2+, Ascl1*+, *Ret*+) containing presumptive sympathetic (*Isl1+*) and ENS (*Isl1-* ) cells (Figure 1G). The remaining *Sox10*+ clusters lacked unique markers and likely represented progenitor or glial cells; these were annotated as NCC1-NCC7 (Figure 1E). Amongst neuron-enriched clusters, cluster 4 displayed a nodose ganglion identity (*Phox2a+, Phox2b+, Tbx1+, Tfap2b-*). Cluster 8 and 12 expressed a DRG profile (*Isl1+, Neurod1+, Six1+, Pou4f1+*), with cluster 12 representing an immature state (higher *Sox10* and *Foxd3* expression) (Figure 1E; Supplementary Figure 2C, D).

To further substantiate cluster identities, we addressed the Hox gene expression patterns (Supplementary Figure 2E, F). Consistent with a cranial origin, the cluster identified as nodose ganglion exhibited robust anterior Hox gene expression (e.g. *Hoxa1-4*, *Hoxb1-3*). ENS and parts of the sympathetic ganglia originate from vagal neural crest, and cluster 6 expressed a characteristic suite of *Hoxd3-4* and *Hoxb6* (*14*). NCC3 lacked Hox gene expression, indicating a rostral cranial identity, while the Hox gene pattern of NCC5 suggested they are progenitors/glia in the nodose ganglion. Posterior Hox gene expression became more pronounced at E10.5, particularly within DRG clusters and several NCC clusters, indicative of the anterior-to-posterior emergence of neural crest derivatives. Together, the data demonstrate that lentiviral injection at E7.5 targets primordia of gut-innervating ganglia, including ENS, sympathetic ganglia, nodose ganglia and DRG (Figure 1D).

To confirm that gut-innervating ganglia are persistently labelled, we performed immunohistochemical analysis of E13.5 DRG and E18.5 gut, in embryos injected at E7.5 with lentivirus carrying H2B-GFP. There was robust transduction of the DRG, with efficient labelling of both NeuN+ neurons and SOX2+ progenitors/glia from thoracic to sacral levels (65-83%) (Figure 1H-J). The ENS was uniformly transduced across gut regions, with an average of 51% HuC/D+ neurons and 44% SOX10+ progenitors/glia (Figure 1K-M; Supplementary Figure 3A-D). In addition to ENS labelling, we observed GFP across all gut layers, including in α-SMA+ smooth muscle cells and EpCAM+ epithelial cells (Figure 1N; Supplementary Figure 3E). This indicates that viral particles in amniotic fluid not only transduce the neural plate (ectoderm) but are also in contact with endo- and mesodermal germ cells at E7.5 (Supplementary Figure 4A, B). Indeed, injections during an early developmental window were crucial for ENS targeting, as injections at E8.0-8.5 failed to label both the ENS and gut epithelium, with only sporadic labeling of mesenchymal cells observed (Supplementary Figure 4C, D).

Together, these data demonstrate that E7.5 amniotic cavity lentiviral injection enables widespread and stable transduction of the GI-tract, encompassing extrinsic gut-innervating neuronal populations, the ENS, and non-ENS gut cell types.

### *In utero* transduction targets all major cell types in the developing gut

As we discovered that injections not only target the ENS but the whole gut wall, we reasoned that this approach could be leveraged to determine lineage relationships between gut cell types derived from all three germ layers. Moreover, by capturing different segments of the gut, conclusions could be drawn about the timing of anterior-posterior regionalization. To simultaneously profile the transcriptional identity (cell type) and lineage history (clonality) of single cells, we applied TREX (11, 12) (Figure 2A, B). TREX relies on *in utero* injection of a high-diversity lentiviral library encoding a 30N (random nucleotides) barcode and tdTomato resulting in genomic integration into dividing progenitors, thereby allowing stable inheritance of the construct by all descendant cells. The expression of the barcode under a synthetic ubiquitous promoter allows its detection by scRNA-seq, enabling reconstruction of clonal relationships among captured cells based on shared barcodes (CloneIDs).

**Figure 2.**
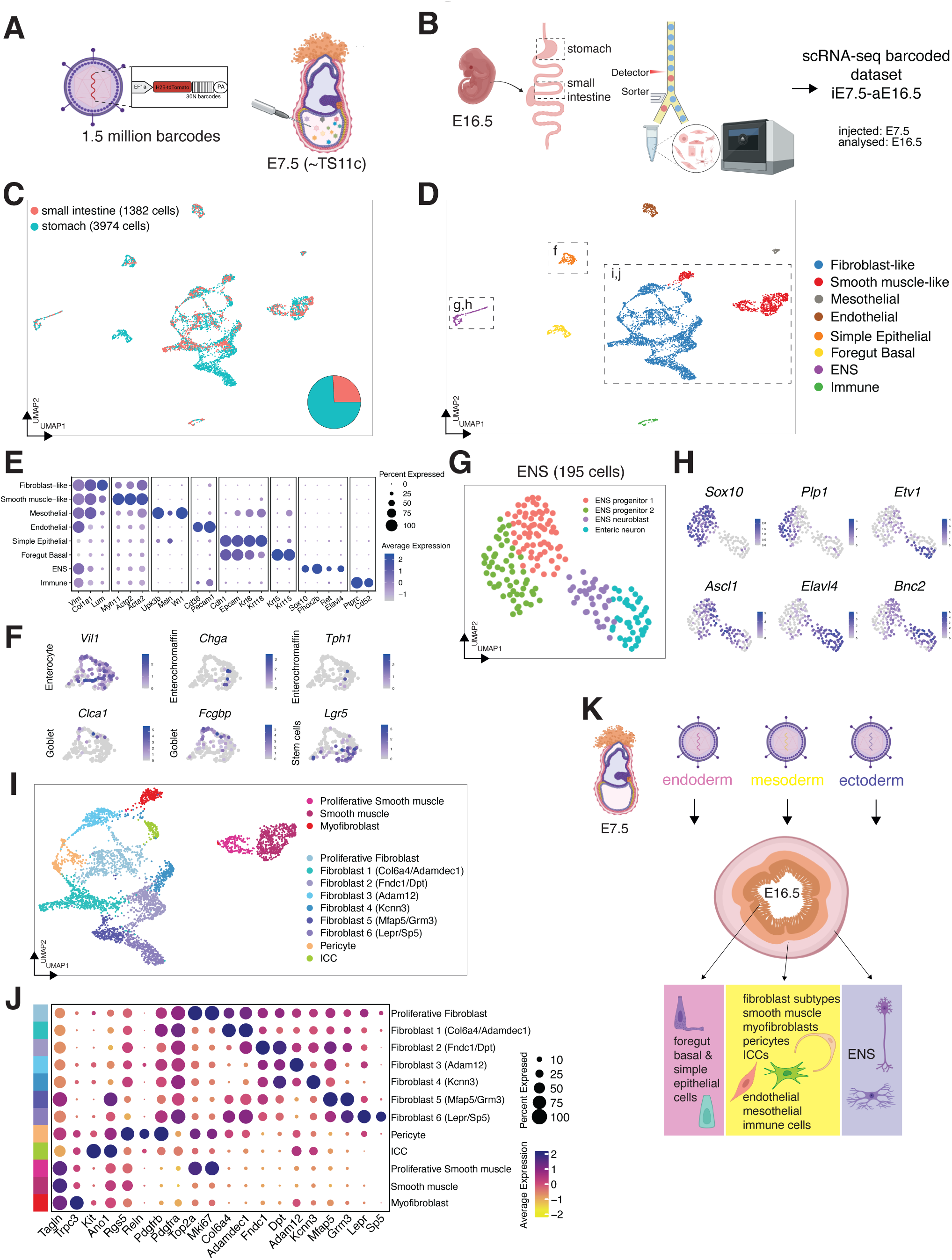
*In utero* transduction at E7. 5 targets diverse cell types in the developing gut. (A) Schematic of injection of lentiviral barcode library into the E7.5 amniotic cavity. (B) Experimental workflow to generate the iE7.5-aE16.5 dataset consisting of TOM+ cells, flow sorted from two gut regions. (C) UMAP plot of high-quality gut cells colored by tissue of origin. Pie chart indicates the proportion of cells from each region. (D) UMAP plot showing 8 major clusters, annotated based on established marker gene expression. (E) Dot plot displaying expression of canonical marker genes used to define major clusters. (F) Feature plots displaying marker gene expression of epithelial cell types. (G) Reprocessed ENS cells revealing four clusters. (H) Feature plots showing expression of selected markers for ENS states and branches. (I) UMAP plot showing subclusters within fibroblast-like and smooth muscle-like populations. (J) Dot plot showing marker gene expression distinguishing fibroblast-like and smooth muscle-like subclusters. (K) Schematic drawing indicating the diverse gut cell types targeted by *in utero* nano-injection, showing that all three germ layers are efficiently transduced. Part of 2A, B and K were created with BioRender.com. (E, J) Dot size represents the percentage of cells expressing the gene, and color indicates relative mean expression.

We injected the tdTomato-30N barcode lentiviral library into the amniotic cavity of E7.5 embryos (Figure 2A) and analyzed at E16.5 (dataset thus called iE7.5-aE16.5). TOM+ cells were sorted from stomach and small intestine by flow cytometry and sequenced using the 10x Chromium platform (Figure 2B). After removing low-quality cells and potential doublets, 5,356 cells were retained for analysis, comprising 3,974 cells from the stomach and 1,382 cells from the small intestine (Figure 2C; Supplementary Figure 5A-C). Cells from both regions were integrated, resulting in eight major clusters defined by canonical lineage markers, corresponding to fibroblast-like cells (*Vim+, Col1a1+, Lum*+), smooth muscle-like cells (*Myh11+, Actg2+, Acta2+*), endothelial cells (*Cd36+, Pecam1+*), foregut basal cells (*Cdh1+, Epcam+, Krt5+, Krt15+)*, simple epithelial cells (*Cdh1+, Epcam+, Krt8+, Krt18+*), ENS (*Sox10+, Phox2b+, Ret+, Elavl4+*), immune cells (*Cd52+, Ptprc+*) and mesothelial cells (*Upk3b+, Msln+, Wt1+*) (Figure 2D, E; Supplementary Figure 5D). Further analysis of the simple epithelial cluster revealed predominantly enterocytes (*Vil1+*), along with smaller populations of enterochromaffin cells (*Chga+, Tph1+*), goblet cells (*Clca1+, Fcgbp+*), and intestinal stem cells (*Lgr5+*) (Figure 2F). The 195 ENS cells formed four transcriptionally distinct clusters representing progenitors, neuroblasts, and neurons (Figure 2G). Notably, previously discovered neuronal branches characterized by *Etv1* (Branch A) and *Bnc2* (Branch B) were captured (Figure 2H) (15).

Subclustering of the smooth muscle-like cluster revealed a contractile myofibroblast-like state (*Trpc3+, Tagln+*) and a proliferative cluster (Figure 2I, J; Supplementary Figure 5D, E). Subclustering of the fibroblast-like cluster uncovered additional mesenchymal heterogeneity, including Interstitial cells of Cajal (ICC) (*Kit+, Ano1+*), pericytes (*Rgs5+, Reln+, Pdgfrb+, Pdgfr*a-) and seven transcriptionally distinct fibroblast subtypes (Figure 21, J). These included proliferative fibroblasts (*Top2a+, Mki67+*), Fibroblast 1 (*Col6a4+, Adamdec1+*), Fibroblast 2 (*Fndc1+, Dpt+*), Fibroblast 3 (*Adam12+*), Fibroblast 4 (*Kcnn3+*), Fibroblast 5 (*Mfap5+, Grm3+*) and Fibroblast 6 (*Lepr+, Sp5+*) (Supplementary Figure 5D-H). The proliferative fibroblast cluster likely comprised a mixture of all fibroblast subtypes. Collectively, our results demonstrate that *in utero* lentiviral transduction effectively transduces progenitors of all major gut cell types, encompassing derivatives of the three germ layers (Figure 2K).

### Barcoded *in utero* transduction reveals clonal lineage structures in the developing gut

The comprehensive capture of diverse gut cell types provided us excellent conditions to reveal detailed lineage relationships. After barcode extraction, 3,959 cells (73.9%) retained CloneIDs (Figure 3A, B; Supplementary Figure 6A-C). Cells with or without CloneIDs showed comparable quality metrics and similar CloneID recovery rates were observed across all cell types, indicating unbiased barcode dropout (Supplementary Figure 6D-G). Most CloneIDs consisted of a single barcode, indicating single lentiviral particle transductions, while up to 4 barcodes were detected in a few cells (0.42% of clones) (Supplementary Figure 6H).

**Figure 3.**
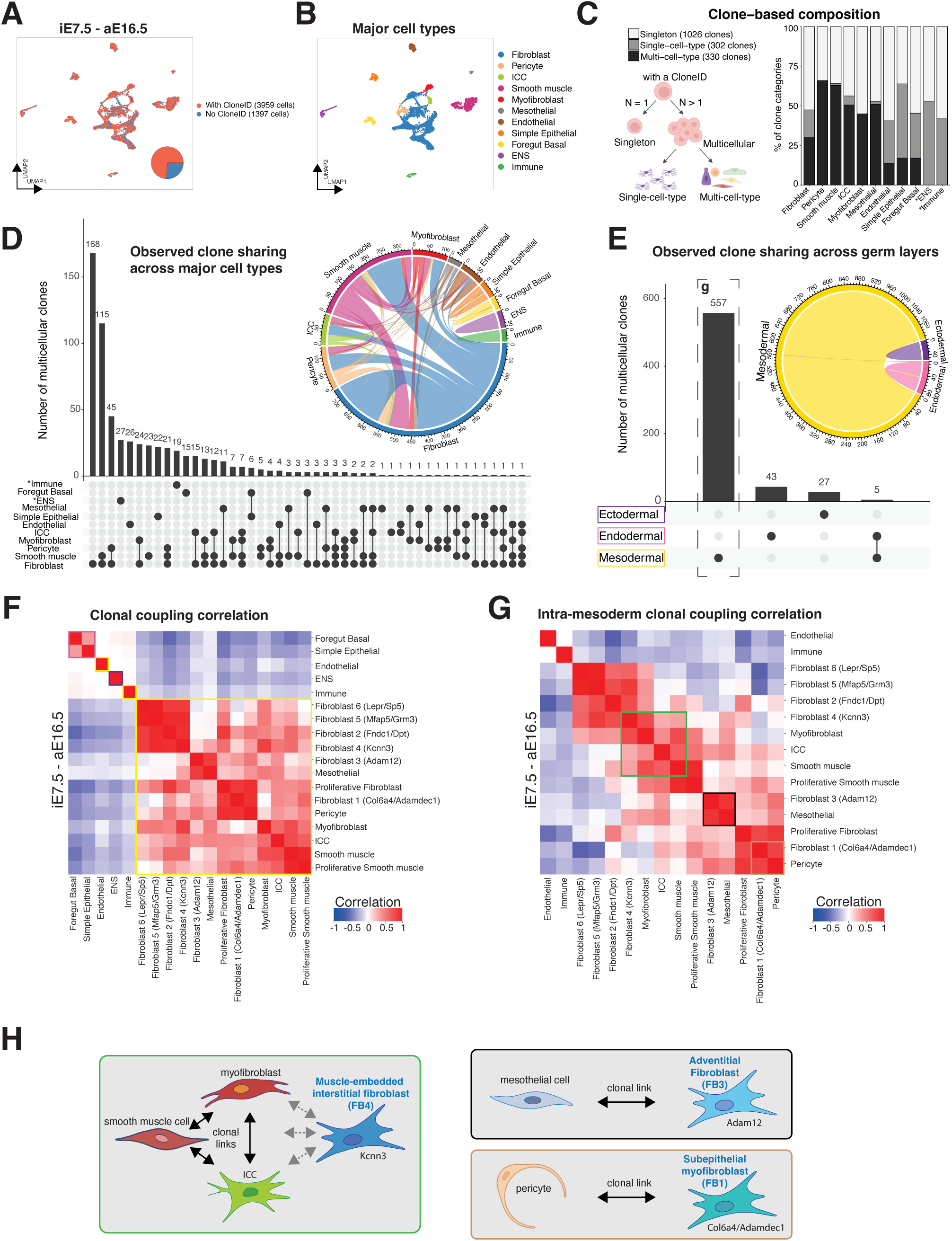
Lentiviral barcoding at E7.5 resolves clonal lineage relations in the developing gut. (A) UMAP plot highlighting proportions of cells with CloneIDs in the iE7.5-aE16.5 dataset. (B) UMAP plot showing 11 major gut cell types after merging subclusters. (C) Stacked bar plot showing distribution of clone categories across major cell types. Schematic drawing describing clonal types (right). (D) UpSet plot showing frequencies of clones with different composition of major cell types. (E) UpSet plot showing minimal clone sharing across germ layers. Inset chord diagrams in (D, E) illustrate the extent and direction of clonal sharing, with ribbon width proportional to the number of shared clones. (F, G) Heatmaps showing the correlation matrix of clonal coupling z-scores, computed as deviations of observed clone sharing relative to randomized data, for refined gut cell types (F) and restricted to mesoderm-derived cell types (G). Colored boxes denote germ-layer origin (pink: endoderm; purple: ectoderm; yellow: mesoderm), or lineage-coupled mesenchymal populations (green: Fibroblast 4 with myofibroblasts, ICC and smooth muscle cells; black: Fibroblast 3 with mesothelial cells; brown: Fibroblast 1 with pericytes). (G) Schematic summary of lineage relationships amongst mesenchymal cell types inferred from the clonal coupling analysis. H and left panel of 3C were created with Biorender.com.

In total, 1,658 clones were reconstructed and classified into three categories: singletons (1,026 clones), single-cell-type clones (302 clones) and multi-cell-type clones (330 clones) (Figure 3C). Multicellular clones contained on average 4.6 cells and reached a maximum of 31 cells (Supplementary Figure 6I). Systematic assessment of multi-cellular clonal compositions per major cell type showed that pericytes, ICCs, smooth muscle and mesothelial cells almost exclusively consisted of multi-cell-type clones (Figure 3C), suggesting substantial lineage diversification. Instead, ENS and immune cells exhibited only single-cell-type clones, reflecting early lineage restriction of their respective precursors. Clone sharing was predominantly found between mesenchymal cell types in the gut, consistent with their shared mesodermal origin (Figure 3D). However, endothelial and mesothelial cells did not share clones with each other despite sharing clones with other mesenchymal cell types, indicating that their progenitor allocation is already distinct at E7.5 (Figure 3D). Cells arising from different germ layers rarely shared clones; only five clones spanned endodermal and mesodermal lineages, likely reflecting residual pluripotent mesendoderm cells (16) at the time of injection (Figure 3E).

To systematically resolve lineage relationships among refined cell types, we performed clonal coupling analysis by calculating z-scores for deviations in observed clone sharing relative to randomized data. Hierarchical clustering of Pearson correlations between z-scores segregated cell types into broad groups corresponding to germ-layer origin and revealed intricate clonal coupling relations between mesoderm-derived populations (Figure 3F, G, Supplementary Figure 6J, K). Notably, ICCs showed strong clonal coupling with smooth muscle cells and myofibroblasts (Figure 3F-H). This is consistent with previous lineage-tracing studies showing that ICCs and myofibroblasts arise from common precursors and that ICCs are not fibroblasts (17). However, ICCs, myofibroblasts and smooth muscle cells also showed clonal coupling with Fibroblast 4 (*Kcnn3*+). This fibroblast likely corresponds to *Pdfgra*+ cells of the SIP (smooth muscle cells, ICCs, and PDGFRa (18, 19)), recently renamed muscle-embedded interstitial fibroblasts; MIFs (20)) (Supplementary Figure 5E, F). Fibroblast 3 (*Adam12+*) showed enriched clonal coupling with mesothelial cells (Figure 3F-H), suggesting an adventitial fibroblast (AF) phenotype (21), further supported by expression of *Wt1* and *Pi16* (Supplementary Figure 5E, G). Fibroblast 1 (*Col6a4+, Adamdec1+*) displayed clonal coupling with pericytes, consistent with the reported localization of *Adamdec1*+ myofibroblasts in the subepithelial space (SEMFs) (Figure 3F-H; Supplementary Figure 5E, H). Notably, clonal coupling analysis thus segregated fibroblast populations into putative contractile, adventitial and perivascular trajectories which was less apparent from transcriptome-based clustering alone (Figure 3H; Supplementary Figure 6L).

Clonal coupling patterns among mesoderm-derived populations indicate that mesenchymal lineages diverge early into fate-biased progenitor pools that underpin coordinated tissue patterning during gut morphogenesis (Figure 3H). Collectively, we demonstrate that single-cell lentiviral barcoding at E7.5 recapitulates canonical developmental hierarchies and uncovers early cell fate diversification within the gut, corroborating known lineage relationships while revealing previously unrecognized trajectories contributing to gut organogenesis.

### Nano-injection at early E7.5 improves ENS targeting while revealing residual epiblasts

Attempting to enrich for ENS targeting, we performed another nano-injection of the N30 barcode lentiviral library aimed at an earlier developmental stage (E7.5^Early^, ∼TS11b; Supplementary Figure 4A, B), reasoning that primordia of neural crest could be physically more accessible and remain for a longer time before delamination. To gain further information about gut regionalization, we collected stomach, small intestine, and large intestine at E16.5 to generate the iE7.5^Early^-aE16.5 dataset (Figure 4A). Data was processed similarly as in the previous experiment, and we recovered the same major cell types (Figure 4B; Supplementary Figure 7A-E). Notably, the earlier targeting increased the proportion of ENS cells by 7.7-fold (Figure 4C), making ENS the most abundant cell population, and corroborating greater lentiviral accessibility to neural crest primordia at the stage of injection. Higher-resolution clustering further resolved subpopulations within immune and ENS lineages, revealing myeloid (*Lyz2*+, *Mrc1*+, *Cd68*+) and lymphoid (*Trbc1/2*+, *Cd4*+) subpopulations, lymphatic signatures (*Lyve1*+, *Pecam1*+), refined ENS neuron divisions, and Schwann cell precursors (SCPs) (Figure 4D, Supplementary Figure 7F, G).

**Figure 4.**
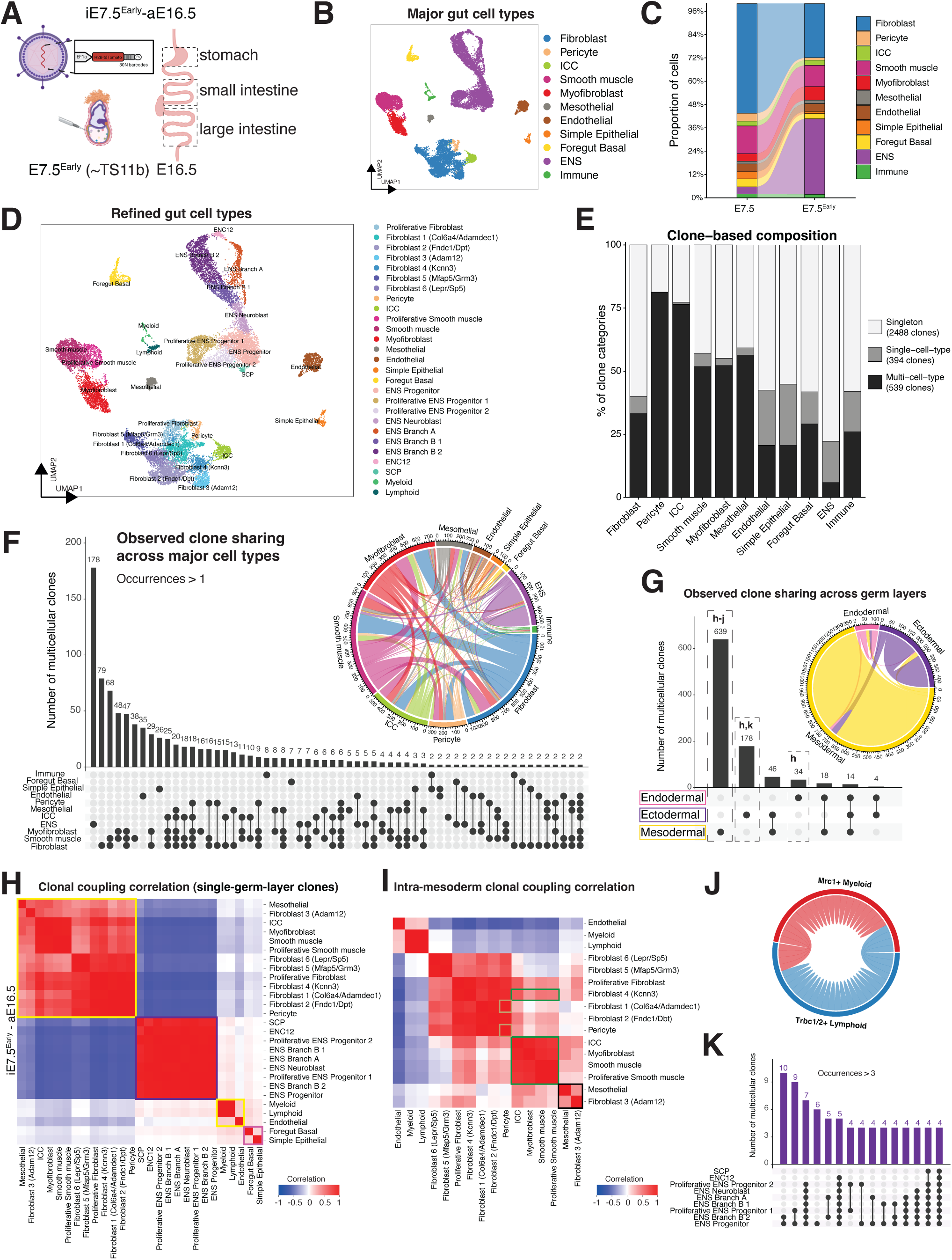
*In utero* transduction at early E7.5 improves ENS targeting but uncovers presence of epiblast cells. (A) Schematic diagram showing the generation of the iE7.5^Early^-aE16.5 dataset consisting of three gut regions. Created with BioRender.com. (B) UMAP plot showing major cell types. (C) Alluvial plot showing proportions of major cell types in the iE7.5-aE16.5 and iE7.5^Early^-aE16.5 datasets. (D) UMAP plot showing refined gut cell type annotations in the iE7.5^Early^-aE16.5 dataset. (E) Stacked bar plot showing distribution of clone categories across major cell types identified in the iE7.5^Early^-aE16.5 dataset. (F) UpSet plot showing frequencies of clones with different composition of major cell types. (G) UpSet plot showing notable clone sharing across germ layers. Inset chord diagrams in (F, G) illustrate the extent and direction of clonal sharing, with ribbon width proportional to the number of shared clones. (H, I) Heatmaps showing the correlation matrix of clonal coupling z-scores, computed as deviations of observed clone sharing relative to randomized data, for refined gut cell types (H) and restricted to mesoderm-derived cell types (I). Colored boxes denote germ-layer origin (pink: endoderm; purple: ectoderm; yellow: mesoderm), or lineage-coupled mesenchymal populations (green: Fibroblast 4 with myofibroblasts, ICC and smooth muscle cells; black: Fibroblast 3 with mesothelial cells; brown: Fibroblast 1 with pericytes). (J) Chord diagram showing self- and cross-linking between Mrc1+ myeloid and Trbc1/2+ lymphoid cells. (K) UpSet plot showing co-occurrence of ENS cell states/types in the same clone. Note that Branch B1/2 are not exact equivalents of Branch B1/2 defined in Morarach et al, 2021 (15).

CloneIDs could be assigned to 77.8% of the 21,749 cells (Supplementary Figure 8A-E), yielding 3,421 clones comprising 2,488 singletons, 394 single-cell-type clones, and 539 multi-cell-type clones (Figure 4E). The average and maximum size of multicellular clones were larger in the E7.5^Early^-aE16.5 dataset (Supplementary Figure 8F). Consistent with the first experiment, mesenchymal populations comprised mainly of multi-cell-type clones, whereas ENS, immune and epithelial populations were more lineage-restricted (Figure 4E). Although most multicellular clones were composed of a single germ layer, a notable fraction (∼9%) of clones spanned more than one germ layer derivative (∼1.5% spanned all three germ layers; 14 clones) (Figure 4F, G). These cross-germ-layer clones suggest the presence of pluripotent epiblast cells at E7.5^Early^, in agreement with recent high-throughput spatial transcriptome analyses of murine gastrula (22).

The presence of cross-germ-layer clones in the iE7.5^Early^-aE16.5 dataset complicated lineage relationship analysis between refined cell types, but we reasoned that analysis using germ-layer pure clones would still be relevant. Clonal coupling analysis of pure clones (851 clones) recapitulated patterns observed in the iE7.5-aE16.5 dataset (compare Figures 3F with 4H; Supplementary Figure 8G). We next restricted analysis to pure mesoderm clones (639 clones) (Figure 4I; Supplementary Figure 8H). As previously observed (Figure 3G, H), ICCs, myofibroblasts and Fibroblast 4 (putative MIFs) featured stronger clonal coupling, as did mesothelial cells and Fibroblast 3 (putative AFs) (Figure 4I). Although Fibroblast 1 (putative SEMFs) was not adjacent to pericytes in the correlation heatmap, it exhibited the strongest clonal pericyte coupling of all fibroblasts (Figure 4I; Supplementary Figure 8H). Within the immune population, *Mrc1*+ myeloid and *Trbc1/2*+ lymphoid cells shared only few clones, indicating early lineage specification prior to gut colonization (Figure 4J). The large ENS dataset allowed detailed clonal analysis of ENS subclusters, revealing that many clones consisted of multiple ENS states/types (Figure 4K). Notably, SCPs showed clonal coupling with all ENS subtypes, however less so with the more mature neuron clusters ENC12 and Branch B2 which may differentiate before SCP arrival to the gut (Supplementary Figure 8G). Coupling was overall strong between Branch A and B neurons (Figure 4K; Supplementary Figure 8G, grey box), suggesting no predetermination of either Branch in agreement with an earlier lineage-tracing study (23).

Together, our results from the E7.5^Early^ injection highlights the importance of precise embryonic staging for enriched targeting of specific lineage precursors. The optimal window for ENS targeting appears to coincide with early E7.5 (∼TS11b stage), when some epiblasts cells are still present. This stage is therefore advantageous for efficient and comprehensive ENS targeting, while posing challenges for lineage analysis.

### Clonal sharing between gut segments reveals lineage-specific timing of regional identity acquisition

Having resolved lineage relationships between and within gut cell types, we next examined the regional dispersion of clones across the upper and lower GI tract. In the iE7.5-aE16.5 dataset, 21 clones linking the stomach and small intestine were identified. Of these, 18 clones (85.7%) consisted exclusively of ENS or immune cells, accounting for 25.9% of ENS and 57.9% of immune clones (Figure 5A-C). These regional dispersions are consistent with the extrinsic origins and migratory behavior of these lineages: the ENS arises from neural crest precursor cells that migrate along the entire gut axis (8), while the first wave of embryonic immune cells originates from the yolk sac and subsequently colonizes the gut (24, 25). By contrast, clones corresponding to other cell types were predominantly restricted to a single gut segment, suggesting that most endoderm- and mesoderm-derived progenitors had acquired their regional identities by E7.5 (∼TS11c).

**Figure 5.**
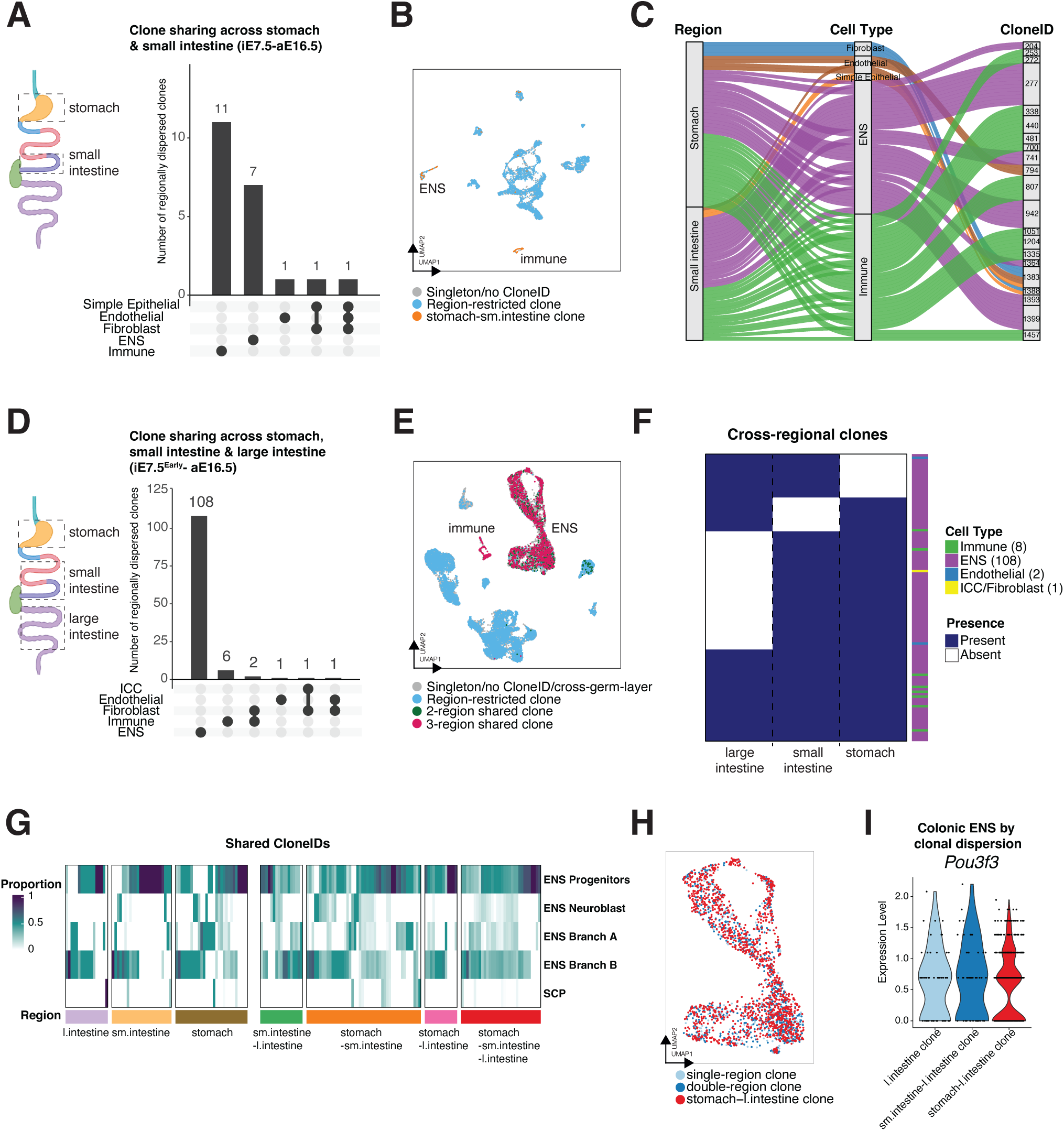
Regional clone sharing reveals long-range ENS migration and early regional patterning of mesenchymal populations along the GI tract. (A) UpSet plot showing co-occurrence of major cell types in cross-region clones in the iE7.5-aE16.5 dataset. (B) UMAP plot highlighting cells in cross-region and local clones. (C) Alluvial plot illustrating relationship between tissue region, cell type, and CloneID of the cross-region clones in (A). (D) UpSet plot showing co-occurrence of major cell types in the cross-region clones in the iE7.5^Early^-aE16.5 dataset (restricted to single-germ-layer clones). (E) UMAP plot highlighting cells in cross-region and local clones. (F) Heatmap showing the distribution of clones across gut regions in the iE7.5-aE16.5 dataset, with binary indication of presence or absence. Colored bars denote cell type identities of clones. (G) Heatmap showing clonal composition of ENS cell states/types for cross-region or local clones, revealing no apparent bias. (H) UMAP plot highlighting the differentiation state of clones with different dispersion. (I) Violin plot showing comparable levels of *Pou3f3* expression in colonic ENS cells that are local or share clones with small intestine and/or stomach. Parts of 5A and D were created with Biorender.com.

Considering single-germ-layer clones in the iE7.5^Early^-aE16.5 dataset, we identified 123 region-spanning clones, again mainly from ENS and immune lineages, accounting for 60.7% of ENS and 66.7% of total immune clones (Figure 5D-F). Despite their abundance, epithelial or fibroblast clones did not span regions, indicating that their progenitors are already regionally patterned also at slightly earlier gastrula stages. We next examined the ENS subtype composition within cross-regional and local clones and found them to be similar. Notably, both Branch A and B neurons were present in local and cross-regional clones, supporting the innate stochastic differentiation program in the ENS (Figure 5G). A distinguishing feature of colonic ENS is its expression of *Pou3f3* (26), while it has not been shown if expression is induced in vagally derived enteric *Pou3f3*- precursors as they enter the large intestine. Our barcoding experiment allowed us to clonally link stomach-resident ENS (derived exclusively from the vagal neural crest) to *Pou3f3*⁺ ENS cells in the colon. We found that stomach/large intestine clones distributed randomly amongst ENS differentiation stages and neuron identities (Figure 5H). Cells within large intestine-only clones and stomach/large intestine clones showed similar expression frequency and level of *Pou3f3* (Figure 5I). Together, these findings support that colonic ENS cells derived from vagal neural crest acquire their colon-specific identity by responding to local cues.

### Identification of cell type-specific regional gene expression differences

Beyond lineage relationships, our regionally resolved scRNA-seq datasets enabled analysis of transcriptional similarities and differences across GI regions at E16.5. Integration of the two datasets showed seamless merging (Supplementary Figure 9A, B), indicating consistent targeting of comparable progenitor populations. Most cell types were detected across all regions except for stomach-specific foregut basal epithelia, which expressed distinct marker genes (Supplementary Figure 9C, D). Small intestine epithelium was instead enriched for *Cps1*, whereas posterior Hox genes marked large intestine epithelium (Supplementary Figure 9D). Region-specific expression patterns were also observed in mesoderm-derived populations: *Barx1* and *Nr2f1* were restricted to the stomach, while *Pitx1* was detected in both stomach and large intestine (Supplementary Figure 9E). *Nkx2.3* and *Mab21l2* were confined to intestinal mesenchyme. Within ectoderm-derived tissue (ENS), *Hoxb5*/*Hoxa4* were enriched across intestinal regions, whereas *Hoxc4* was limited to the small intestine (Supplementary Figure 9F).

Focused analysis of fibroblast subtypes revealed that Fibroblasts 5 and 6 were enriched in the stomach compared with the intestines (Supplementary Figure 9G, H). Within these populations, *Grm3* and *Sp5* showed stomach-specific expression (Supplementary Figure 9I), whereas Fibroblast 6 uniquely expressed *Thy1* (Cd90), a marker associated with crypt fibroblasts (Supplementary Figure 9I). *Wnt6*, *Cpne8* and *Ism1* in Fibroblast 6 further supported a fibroblast type with high signaling capacity, consistent with a crypt/submucosal-associated niche (Supplementary Figure 9I). These differences may reflect either a greater abundance of Fibroblast 5 and 6 in the stomach or their earlier differentiation in anterior gut regions.

In summary, our collective E16.5 scRNA-seq data uncovered regionally distributed cell types, expression patterns and fibroblast states in the embryonic gut, which may contribute to establishing the functional specialization of individual GI regions.

### Cre-dependent lentivirus nano-injection enables cell type-specific targeting

Our data show that progenitors of essentially all gut cell types are transduced by *in utero* lentiviral nano-injection at E7.5. For applications requiring cell type-specific targeting, this approach therefore needs adaptation. Previous studies have demonstrated that mini-promoters can confer cell type-specificity in the developing CNS (10). However, no ENS-specific promoters have yet been described. Instead, Cre-driver mouse lines have been used to label all or subsets of ENS cell populations.

We previously showed that Baf53b-Cre mice specifically label neurons in the juvenile ENS (15). To investigate their utility during embryonic development, Baf53b-Cre; R26-tdTomato embryos were analyzed at two stages (Supplementary Figure 10A, B). Reporter expression was restricted to HuC/D+ neurons and absent from SOX10+ progenitor/glial cells. We found that ∼50% of HuC/D+ neurons expressed TOM at E14.5, increasing to ∼90% at E18.5. Baf53b-Cre activity is thus confined to differentiating neurons, with a slight delay relative to HuC/D expression.

To achieve conditional targeting with *in utero* nano-injection, we designed a lentiviral vector expressing an inverted double-floxed EGFP reporter, Lenti-DIO-EGFP (Figure 6A, Supplementary Figure 1A). While AC injection of non-conditional LV-H2B-GFP resulted in GFP labelling of HuC/D+ neurons, GFP was additionally observed in non-ENS cells as previously discussed (Figure 6B; 1N). In contrast, AC injection of Lenti*-*DIO-EGFP into Baf53b-Cre embryos resulted in GFP expression restricted to HuC/D+ neurons, labelling ∼4-6% at E18.5 (Figure 6B, C). In conclusion, *in utero* nano-injection of conditional lentiviral vectors enables precise targeting of specific gut cell types, as demonstrated by Baf53b-Cre-dependent lentiviral reporter expression in enteric neurons.

**Figure 6.**
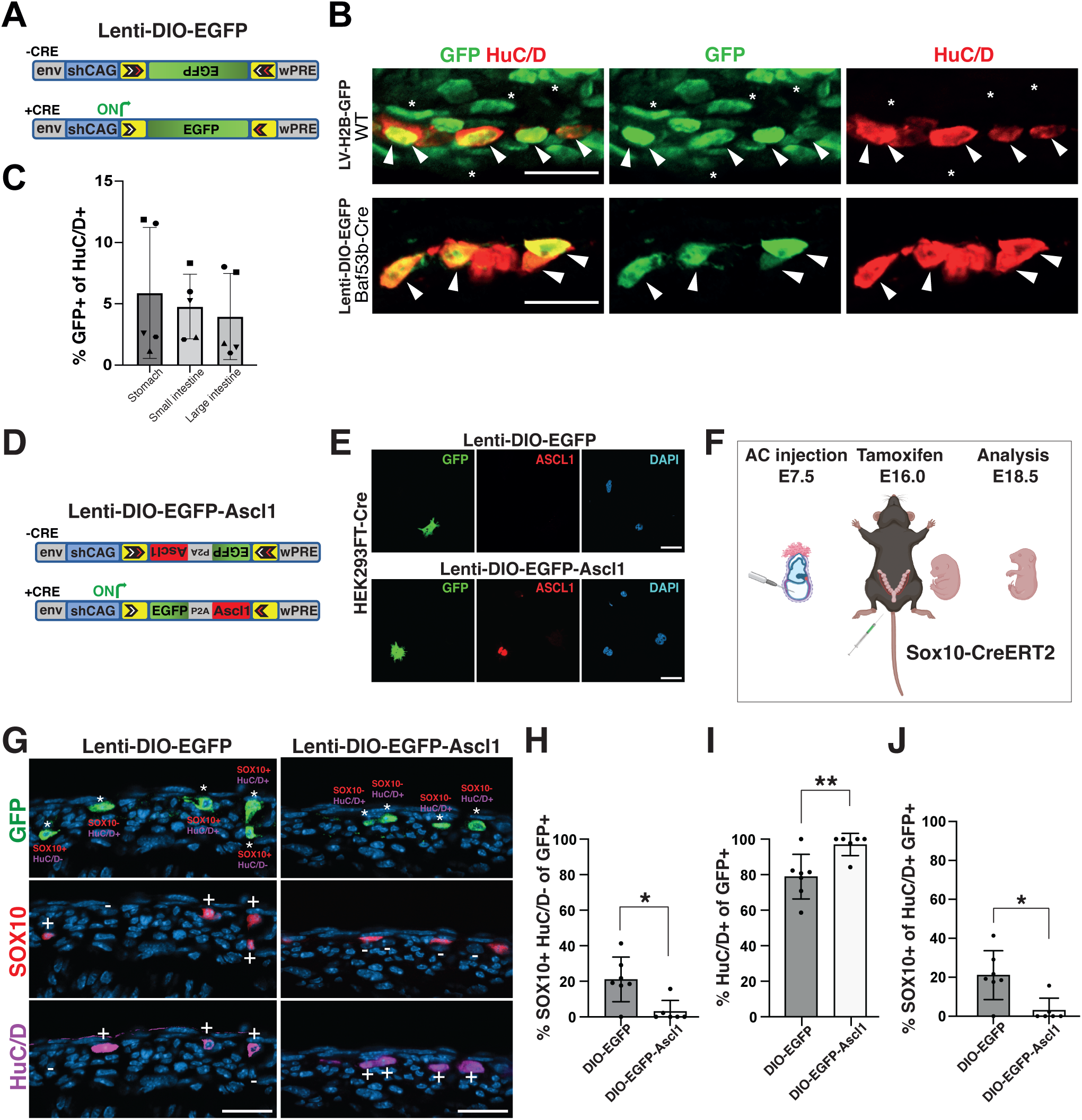
Temporal overexpression of Ascl1 in Sox10+ cells promotes neuronal differentiation. (A) Schematic overview of the Lenti-DIO-EGFP vector, before and after Cre-mediated gene inversion. (B) Representative immunohistochemistry images of intestine from wild-type mice injected with LV-H2B-GFP showing GFP expression in both HuC/D- cells (asterisks) and HuC/D+ (arrowheads), and Baf53b-Cre mice injected with Lenti-DIO-EGFP showing GFP expression restricted to HuC/D+ cells (arrowheads). (C) Bar graph showing the percentage of HuC/D+ cells expressing GFP across gut regions in Baf53b-Cre embryos injected with Lenti-DIO-EGFP. (n=5 mice). Bars represent mean ± SD. (D) Schematic drawing of the Lenti-DIO-EGFP-Ascl1 construct with and without Cre-mediated gene inversion. (E) Representative immunohistochemistry images showing validation of transgene expression in Cre-expressing HEK293FT cells. (F) Schematic representation of experimental procedure for *in vivo* overexpression of EGFP-Ascl1 or EGFP in ENS progenitors. Part was created with BioRender.com. (G) Representative confocal images of E18.5 small intestine transduced with DIO-EGFP or DIO-Ascl1-EGFP vectors showing SOX10 and HuC/D expression patterns in GFP+ cells. asterisks denote EGFP+ cells and texts indicate their status of expression, with + or – signs denoting positive or negative expression. (H-J) Bar graphs showing the proportions of GFP+ cells with SOX10 expression without HuC/D (H), GFP+ cells with HuC/D expression (I), and GFP+ HuC/D+ cells expressing SOX10 (J), in Baf53b-Cre embryos injected with either DIO-Ascl1-EGFP or Lenti-DIO-EGFP. (H)*p=0.0136; (I)**p= 0.0086, and (J)* p= 0.011. n=6-7 mice per condition. SI: small intestine; AC: amniotic cavity; WT: wild-type. Scale bars: 20µm.

### Timed overexpression of Ascl1 promotes neuronal differentiation of ENS progenitors

Having validated that lentiviral DIO-EGFP expression can be controlled by Cre, we next sought to use this approach to drive timed gene expression in ENS progenitor cells, which are more responsive to developmental cues. Sox10-CreERT2 mice mediate efficient, temporally controlled recombination in ENS progenitors (13, 14, 23). To assess the efficiency and kinetics of Cre-dependent recombination, tamoxifen was administered to Sox10-CreERT2 females mated with R26-EYFP males at E16.5, and embryos were harvested 26 hours later. Out of SOX10+ cells, ∼50% expressed EYFP and ∼11% of HuC/D+ neurons, indicating efficient labeling of progenitors/glia and a subset of newly differentiated neurons within a short time window (Supplementary Figure 10C-E).

As a proof-of-principle, we sought to overexpress *Ascl1*, a proneural transcription factor essential for enteric neurogenesis, as demonstrated by us and others (27, 28). We generated a Cre-inducible *Lenti-DIO-EGFP-Ascl1* construct, in which *Ascl1* was linked to EGFP via a self-cleaving P2A peptide (Figure 6D, Supplementary Figure 1B). Cre-dependent ASCL1 expression was validated using a HEK293FT-Cre cell line (Figure 6E). We next overexpressed ASCL1 at late embryonic stages using our system, when both glio- and neurogenesis are ongoing, to assess its ability to promote neuronal differentiation. Sox10-CreERT2 embryos were injected at E7.5 with lentivirus carrying either DIO-EGFP as controls or DIO-EGFP-Ascl1, followed by tamoxifen administration at E16.0 and analysis at E18.5 (Figure 6F). As compared to controls, ASCL1 overexpression reduced the EGFP+ progenitors/glia population (SOX10+ HuC/D-) from ∼20% to ∼5% (Figure 6G, H) and increased EGFP+ neuronal population (HuC/D+) from ∼80% to ∼95% (Figure 6G, I). Notably, ∼25% of HuC/D+ EGFP+ cells in controls co-expressed SOX10, indicative of a transitional population, which was reduced to <5% upon ASCL1 overexpression (Figure 6G, J).

Together, these findings demonstrate that ASCL1 overexpression during a gliogenic developmental window is sufficient to bias enteric progenitors toward neuronal differentiation. More broadly, this approach establishes ultrasound-guided *in utero* delivery of Cre-inducible lentiviral constructs as a powerful method for temporally controlled, cell type-specific gene manipulation in developing embryos.

## DISCUSSION

Our understanding of gut development and related diseases has advanced substantially through recent technologies, including scRNA-seq and spatial transcriptomics (15, 26, 29). However, methods to define lineage relationships and interrogate gene function *in vivo* have lagged behind, partly due to the inaccessibility of embryonic gut using traditional targeting methods. Here, we present ultrasound-guided AC nano-injection as an efficient methodology to access major gut cell populations in mouse embryos, including epithelial and mesenchymal populations, the ENS, as well as gut-extrinsic ganglia. Combined with high-diversity lentiviral barcoding, this approach establishes a versatile lineage-tracing platform, and coupled with Cre-dependent constructs, the approach supports temporally controlled, cell type-specific gene manipulation in the developing gut.

Barcode-based lineage tracing combined with scRNA-seq overcomes key limitations of traditional Cre-based lineage tracing approaches as they depend on targeting stem cells defined by single marker genes with often equivocal specificities that risk confounding interpretation. In contrast, the method presented here is unbiased and enables comprehensive reconstruction of lineage relationships. It allowed us to resolve unprecedented details amongst transcriptionally related mesenchymal populations refining their cellular identities, lineage relationships and early development. First, we identified three fibroblast states with distinct clonal relationships to specialized mesenchymal cell types. *Kcnn3*+ fibroblasts – putative contractile fibroblasts (MIFs) – were clonally related to ICCs and myoblast/smooth muscle cells, suggesting shared progenitors. *Adamdec1*+ fibroblasts (putative SEMFs) were linked to pericytes, whereas *Adam12*+ fibroblasts (putative AFs) were clonally related to mesothelial cells. Notably, although previously suggested marker genes were used to annotate fibroblast subtypes, no consensus nomenclature currently exists for gut fibroblasts. Clonal analysis therefore provided important orthogonal support for subtype assignments and supported a developmental logic whereby stromal gut compartments are assembled by a set of fate-biased mesenchymal progenitor populations. Second, by analyzing clonal dispersion across gut regions, we demonstrate that gut mesoderm is already regionally specified along the anterior-posterior axis by E7.5, coinciding with persistent epiblast cells. Third, we present regional diversity amongst mesenchymal cell types, likely reflecting early programs that shape region-specific gut wall composition. Overall, these findings highlight how lineage information complements transcriptomic and anatomical analyses to better define gut stroma development. Increasing evidence shows that different fibroblast subpopulations may have distinct roles in cancer, inflammation, and fibrosis (30), underscoring the importance of better understanding their heterogeneity and development.

By integrating multiple gut regions, we found that region-spanning clones were common in migratory lineages such as ENS and immune cells. Within the ENS, clonal sharing across stomach and colon supports a model in which vagal neural crest-derived progenitors acquire region-specific features, such as colonic *Pou3f3* expression, in response to local cues. In contrast, the restriction of epithelial and mesenchymal clones to single regions indicates that regional identity of these compartments is established incredibly early. Together with the region-specific transcriptional programs observed in mesenchymal subtypes, these findings suggest that developmental patterning of the gut wall emerges from a combination of early regional allocation and local differentiation programs.

Beyond lineage reconstruction, *in utero* nano-injection enables conditional gene manipulation in defined ENS populations. Using Baf53b-Cre and Sox10-CreERT2 mice, we show that Cre-dependent lentiviral cargo can be activated in differentiating enteric neurons or temporally controlled progenitor/glial populations. Proof-of-principle experiments with *Ascl1* demonstrated that the system can be used to test gene function *in vivo* during defined developmental windows. This approach complements emerging fetal atlases and disease genetics studies, which increasingly identify candidate regulators in transient embryonic cell states but often lack efficient *vivo* validation strategies. We believe that the ability to combine clonal analysis with targeted perturbation can help shift descriptive cell atlas data to mechanistic studies of gut development.

Several limitations should also be considered. Our results highlight a strong temporal component of lineage accessibility and commitment. Targeting at E7.5^Early^ enhanced labeling of ENS precursors but also increased cross-germ-layer clones, consistent with presence of epiblast cells. While residual epiblast labeling complicates clonal interpretation, it also provides a functional readout of the developmental window during which germ-layer restriction is being consolidated *in vivo*. Precise embryonic staging is thus essential, and inter-embryo variability can complicate stage estimation, although ultrasound imaging partially mitigates this issue. Tissue dissociation, single-cell capture, and barcode recovery also introduce dropouts that will inevitably reduce clonal reconstruction efficiency. In the gene perturbation setting, conditional targeting efficiencies in the ENS were relatively modest. However, we believe that this mosaicism may in fact be advantageous for studying cell-intrinsic mechanisms given that ENS development depends on coordinated proliferation and long-range migration. Broad perturbation within the ENS may cause large secondary effects at the population level, whereas mosaic manipulation may reveal how genetically altered cells behave within a largely normal developing ENS.

Finally, this work has implications for understanding the developmental origins of GI disease. Gut organogenesis relies on interdependent differentiation of diverse cell populations from all three germ layers, and subtle deviations in lineage allocation, maturation, or regional patterning may have lasting functional consequences (31). This is relevant not only to congenital disorders such as Hirschsprung disease, achalasia, and CIPO, but also to more common conditions in which early developmental alterations may predispose to later dysfunction. Some of these disorders may result from *de novo* mutations, in which early mutations occur in a mosaic fashion (32). It is conceivable that the impact of such mutations would be more predictable with a clearer understanding of refined cell lineages arising in post-gastrula stages. We propose that this platform will provide a scalable strategy for mechanistic testing of candidate disease regulators, and modeling early cellular events that may contribute to GI disorders and enteric neuropathologies.

## ACKNOWLEDGEMENTS

We thank Jonas Frisén and Michael Ratz for designing and sharing the tdTomato 30N barcoded viral plasmids. We also thank Jonas Frisén for access to flow cytometry at Karolinska Institutet. We thank Fernando Lopez-Redondo for laboratory assistance. The authors acknowledge support from the National Genomics Infrastructure in Stockholm funded by Science for Life Laboratory, the Knut and Alice Wallenberg Foundation and the Swedish Research Council. The authors acknowledge the computational resources provided by the National Academic Infrastructure for Supercomputing in Sweden (NAISS), partially funded by the Swedish Research Council through grant agreement no. 2022-06725 and the Centre for Bioinformatics and Biostatistics (CBB) at Karolinska Institutet funded by CIMED Infrastructure and Karolinska Institutet. The imaging was performed at the Biomedicum Imaging Core with support from the Karolinska Institutet. We thank the Viral Vector Facility (VVF) of the Neuroscience Center Zurich (ZNZ) for excellent service, construction and production of Lenti viral plasmids. We thank animal staff from the Department of Comparative Medicine at Karolinska Institutet for help with animals.

U.M. was supported by the Swedish Research Council (Vetenskapsrådet; 2020-01129 and 2022-01570), the European Research Council (divENSify; 101045026), the Knut and Alice Wallenberg Foundation (KAW; 2020.0109), the Strategic Research Area in Neuroscience, the Strategic Research Area in Stem Cell Research and Regenerative Medicine, Karolinska Institutet (KI) Consolidator Grant, and the Brain Foundation (Hjärnfonden; FO2025-0169; FO2023-0130). E.R.A was supported by KAW (2020.0109), KI Consolidator Grant (2-195/2021), the Strategic Research Area in Neuroscience, Hjärnfonden (FO2025-0271), Vetenskapsrådet 3R grants (2024-03113, 2021-03537, 2017-01054), and the European Research Council (LIMITLESS; 101171156). K.P was supported by Marie Sklodowska Curie Award (1011544005; DeSTENy). K.H, J.H and Z.L were supported by a KI Grant for Salary (2-2110;2019-7;2019-3).

