## Supplementary Information for "Resolving cell lineages and gene functions in the developing mouse gastrointestinal tract using *in utero* transduction"

### Supplementary Information for Liu Z, Padmanabhan K *et al* 2026

#### Supplementary Material and Methods

##### Plasmid preparation

Plasmid *LV-H2B-GFP* (Addgene plasmid #25999), *Lenti-DIO-EGFP, Lenti-DIO-EGFP-2A-Ascl1* were transformed into bacteria, which were grown in 100mL Terrific Broth containing 100µg/mL Carbenicillin, on a horizontal shaker for 20-22 hours (h). Bacteria were centrifuged to pellet at 4000x g for 20 minutes (min) at 4°C. For DNA extraction PureLink™ or Macherry Nagel Endotoxin-Free Maxi Plasmid Purification Kit were used and concentrations measured by NanoDrop. All lentiviral constructs are third-generation lentiviral vectors.

##### Production of lentiviruses

High-titer vesicular stomatitis virus G (VSV-G)–pseudotyped lentiviral particles were produced either in-house or by a commercial facility (GEG TECH, France). For in house production, low passage (p4-p6) HEK 293 cell line with stable large T antigen expression (Lenti-XTM 293T cells (Clontech) or HEK293 FT cells (Thermofisher)) were thawed one week prior to virus production and grown at 37°C/5% $CO_2$ in growth medium (DMEM high glucose, GlutaMAX pyruvate, supplemented with 10% fetal bovine serum (FBS) and 1% penicillin/streptomycin (Pen/Strep). To sustain stable large T antigen expression, Geneticin (1%) was included in the growth medium during routine maintenance and propagation but was omitted during viral production and transduction procedures. Viral production was performed by transient transfection of the cells followed by cell culture for up to 60 h post-transfection. Calcium phosphate–based and polymer-based transfection reagents, including polyethyleneimine (PEI) and PolyJet (SignaGen laboratories), were used, both methods yielding high viral titers exceeding $1 \times 10^9$ transducing units (TU)/mL. At 12-16 h post-transfection, the growth medium with serum (Polyjet transfection protocol) or without serum

(Calcium phosphate and PEI transfection protocol) was replaced with fresh growth media. Viral supernatant was collected every 24 h until 60 h post-transfection, with fresh medium replenished at each collection. Approximately 280 ml of pooled culture supernatant was concentrated to a final volume of <4 ml by multiple rounds of centrifugation through 100 kDa molecular weight cutoff centrifugal filter units (Millipore Centricon 70 Plus) at 3,300 × g and 4°C, with individual centrifugation steps lasting 30–60 min.

The concentrated supernatant was transferred to ultracentrifuge tubes, and viral resuspension buffer (VRB; 20 mM Tris-HCl, pH 8.0, 250 mM NaCl, 10 mM $MgCl_2$, 5% sorbitol) was added to a final volume of 4 ml. The mixture was homogenized by vigorous pipetting. To form a sucrose cushion, 500 µl of 20% (w/v) sucrose in PBS was carefully layered at the bottom of each ultracentrifuge tube, creating a distinct phase beneath the viral suspension. Viral particles were pelleted by ultracentrifugation at 45,000 rpm for 1.5 h using an MLS-50 rotor. Following centrifugation, the supernatant was removed, and the viral pellet was resuspended in 20–30 µL of VRB or PBS. Aliquots were stored at −80°C.

**Lentiviral titration**

Lentiviral titers were determined by flow cytometry or quantitative (q)PCR. HEK293FT or NE4C cells were used as indicator cells. Cells were seeded in 6-well plates, and the cell number was determined after 12 h from one well. The remaining wells were transduced with serial dilutions of lentiviral supernatant in infection medium (DMEM high glucose, GlutaMAX pyruvate, growth medium supplemented with 20% FBS and 0.1 mg/mL polybrene). Plates were centrifuged at 1100 × g for 30 min at 37°C (spinoculation). The infection mixture was replaced with fresh growth medium, and cells were incubated at 37°C. 96 h after transduction; cells were trypsinized and analyzed by flow cytometry. Alternatively, GFP-positive cells were quantified by fluorescence microscopy. Infectious titers (infectious units per milliliter, IFU/mL) were calculated using the formula: IFU/mL = (% GFP-positive cells × number of cells at the time of transduction) / (volume of virus × dilution factor). For qPCR-based titration, viral RNA was isolated using the NucleoSpin RNA Virus kit (Macherey-Nagel), and lentiviral genome copies were quantified using the Lenti-X qPCR Titration Kit (Takara Bio) according to the manufacturer's instructions.

**Validation of DIO lentiviral constructs**

Low-passage HEK293FT cells (P4–P6) were cultured at 37°C in 5% $CO_2$ and seeded at $5 \times 10^4$ cells per well in poly-L-lysine–coated 6-well plates upon reaching ~85% confluence. After 24 h, cells were transduced with lentiviral Cre-Puro particles ($1 \times 10^7$ particles per well) in infection medium, followed by centrifugation at 1,100 × g for 30 min to enhance transduction efficiency. Medium was replaced post-centrifugation, and 48 h later, cells underwent puromycin selection (1 µg/mL) for 3 days. Surviving cells were expanded and cryopreserved. Stable Cre-expressing HEK293FT cells were reseeded under identical conditions and transduced with Lenti-DIO-EGFP or Lenti-DIO-EGFP-Ascl1 lentiviral particles ($1 \times 10^7$ particles per well) using the same infection protocol. At 96 h post-transduction, cells were replated onto poly-L-lysine–coated glass coverslips in 24-well plates at $2 \times 10^4$ cells per well. The cells were fixed 24 h later in 4% paraformaldehyde (PFA) for 10 min at room temperature (RT) and permeabilized with 0.3% Triton X-100 for 5 min, followed by immunolabeling for ASCL1 and GFP using standard procedures. Briefly, cells were blocked with phosphate-buffered saline (PBS) containing 0.5% bovine serum albumin (BSA) for 1 h, followed by incubation with primary antibodies for 1 h at RT. Coverslips were washed three times with PBS and incubated with secondary antibodies for 1 h at room temperature. After three additional PBS washes, coverslips were mounted onto glass slides for imaging.

**Ultrasound-guided *in utero* nano-injections**

A detailed description of the procedure has been reported (1, 2). Briefly, pregnant females were anesthetized and placed in the supine position on a heated surgical platform maintained at 37 °C. Analgesia was administered prior to surgery. A minor laparotomy was performed to expose the uterus. At a given time, 2-3 embryos were gently exteriorized through an elastic membrane positioned at the base of a modified Petri dish and immobilized using a columnar support of modeling compound. The dish was filled with PBS, and the amniotic cavities were visualized using a high-frequency ultrasound system (VEVO 2100, VisualSonics) with the probe submerged in PBS. A glass microcapillary needle (outer diameter, ~40 μm) attached to a nanoinjector (Nanoject III; Drummond Scientific) and loaded with viral vector was similarly lowered into the PBS and aligned with the ultrasound probe to permit visualization and targeting of the amniotic cavity. Amniotic cavities were injected with the maximal permissible volume, which was defined according to amniotic cavity size and mouse strain as described below. Embryos were gently returned to the maternal abdominal cavity. The muscular layer was closed with 6-0 Prolene sutures, and the skin was closed using EZ clips.

**Comments on embryonic stage and injection parameters**

High-resolution ultrasound imaging of embryos from CD1 and C57BL/6 mice enabled identification of all corresponding Theiler stages (TS) spanning the E7.0–8.5 developmental window, as defined by eMouseAtlas Project (Supplementary Figure 4A-C). We considered Theiler stages TS11A–C to represent the optimal E7.5 window for ultrasound-guided *in utero* injections (Supplementary Figure 4A, B), while injection at E8 or later failed to efficiently target gut and ENS (Supplementary Figure 4C, D). Strain-specific differences were observed, with CD1 embryos displaying larger amniotic cavities and overall embryonic structures, facilitating *in utero* nano-injection. Embryos at TS11A-C, tolerated volumes of lentiviral particles ranged from 30 to 207 nL, consistent with our earlier studies (1, 2). The more robust CD1 embryos were used for lineage tracing experiments and assessment of labeling efficiency. All Cre driver lines for ENS cell type–specific genetic perturbations were maintained on the C57BL/6 background.

**Immunohistochemistry**

Primary antibodies used: chicken anti-GFP (1:1000; Abcam AB13970); sheep anti-GFP( 1:500; Bio-Rad #4745-1051); mouse anti-NeuN (1:200 Millipore MAB377); goat anti-SOX2 (1:200; Santa Cruz SC-17230); mouse anti-HuC/D (1:300; Molecular Probes A21271); rabbit anti-HuC/D(1:300; Abcam ab184267); human anti-HuC/D (1:100; ANNA-1 clone gift from ); goat anti-SOX10 (1:100-500; R&D AF2864); rabbit anti-SOX10 (1:300; Clone #EPR4007, Abcam AB155279); rabbit anti-αSMA (1:500; Abcam5694); rat anti-E-cadherin (1:200; Biolegend 147301); rabbit anti-ASCL1 (1:500; Abcam ab211327).

Secondary antibodies used: donkey anti-chicken 488 (1:400; Jackson ImmunoResearch 703-545-155); donkey anti-sheep 488 (1:400 ThermoFisher; AB 2534082); donkey anti-rabbit 555 (1:1000; ThermoFisher A31572); goat anti-mouse IgG2b 555 (1:1000, ThermoFisher A21147); donkey anti-goat 647 (1:250, ThermoFisher A21447); donkey anti-Rat 647 (1:250, Invitrogen A48272); donkey anti-human Cy5 (1:250; Jackson Immuno Research 709-165-149) and DAPI (1:1000 Sigma D9542).

**Cell counting and statistical analysis**

Immunofluorescence staining was assessed and images acquired using the Zeiss LSM confocal microscopes in the series LSM710 - LSM980 Airy. Animals from different litters were used for repeat experiments to avoid confounding. Quantifications were performed on confocal images, using ‘cell counter’ plugin in ImageJ or manually. For DRG quantifications 8 sections were used per embryo, 134μm apart and spanning from thoracic to sacral levels. Data were analyzed using GraphPad Prism 9 and presented as mean ± SD. Two-sided student's t-test was used to determine statistical significance.

**Embryonic tissue preparation for single-cell RNA sequencing**

CD1 embryos injected with LV-EF1A-H2B-tdTomato-30N were harvested at E16.5. Stomach and 1/3 of the distal small intestine were harvested from the E7.5 experiment, while the stomach, 2/3 of the small intestine, and the entire large intestine were harvested from the E7.5$^{Early}$ experiment. Tissue was kept in cold DMEM/F12 medium (Invitrogen) on ice during dissection. The mesentery was removed, the intestines were cut into 2–5mm$^2$ pieces and put into a digestion solution (0.75 mg/ml Liberase TH (research grade, Roche), 0.1 mg/ml DNase I (Sigma), 4 U/ml dispase (Corning®, 354235) for stomach, 3 U/ml dispase for small and large intestine, in DMEM/F12) at 37 °C for 15 min for the stomach or 12 min for the small and large intestine with shaking every 3 min. The digestion mixture was replaced with DMEM/F12 medium containing 2% BSA (ThermoFisher) and 5 mM EDTA (Invitrogen). The cells were then manually triturated using three fire-polished Pasteur pipettes with decreasing bore diameters (from ~70% to 10% of the original bore diameter) that were previously coated with 1% BSA solution at RT. The single-cell suspension was filtered through a DMEM/F12-equilibrated 30-μm cell strainer (Miltenyi Biotec) and centrifuged at 200x g for 10 min at 4°C. Cells were resuspended in DPBS (Gibco) containing 1% BSA and then incubated with DRAQ7 Dye (1:200) (Biostatus) for 5 min before sorting. Tomato+ DRAQ7- cells were sorted into a 1.5mL Eppendorf tube coated with 1% BSA using a BD Influx (v7) Cell Sorter or a BD FACSAria Fusion equipped with a 100 μm nozzle and were collected in ice-cold DPBS containing 0.04% BSA. Gating was performed using BD FACS software v. 1.0.0.650 or BD

FACSDiva (v9.0.1) (see examples in Supplementary Figure 1C-H). After sorting, samples were centrifuged at 200x g for 8 min at 4 °C, and supernatants were carefully discarded.

**Single cell RNA-sequencing**

cDNA libraries were prepared according to the manufacturer's instructions, using 10x Chromium Next GEM Single Cell 3' Reagent Kits Version 3.1 Dual Index (10x Genomics, CA, USA). Cell suspensions were adjusted to 500–1,000 cells per μl and added to 10x Chromium RT mix to achieve target cell recovery of between 3,000–10,000 cells per reaction. The libraries were sequenced on Illumina NovaSeq 6000 or XPlus platforms using Novaseq S4 reagent kit; 300 cycles. The reads were trimmed to 28-10-10-90 and demultiplexed on the Data Delivery System at NGI Sweden. The datasets were named according to stage of injection (i) and analysis (a): iE7.5-aE16.5 and iE7.5$^{Early}$-aE16.5.

**Analysis of single cell RNA-sequencing datasets**

***Analysis of E9.5 and E10.5 datasets***

Single cell RNA-sequencing datasets from Lentivirus-injected 9.5 and E10.5 embryos(3) were re-analyzed using Scanpy (v1.11.4). The neural crest cell cluster was subsetted from the full dataset and reprocessed to identify subpopulations. Specifically, counts were renormalized, highly variable genes (HVGs) were recalculated within the subset, and principal component analysis (PCA), neighbourhood graph construction, Uniform Manifold Approximation and Projection (UMAP) embedding, and Leiden clustering were repeated. Subcluster identities were assigned based on differential gene expression analysis using the Wilcoxon rank-sum test (sc.tl.rank_genes_groups) as implemented in Scanpy.

***Single-cell RNA sequencing preprocessing for E16.5 embryos***

A custom reference was generated by combining the mouse reference genome GRCm38 (mm10, version 2020-A) genome and an additional sequence representing tdTomato-N transgene, in which the barcode region is marked with 'N' 30-bp barcode wildcard characters using 10x Genomics CellRanger v5.0.(3, 4) Single-cell sequence FASTQ files were aligned to this customized reference using CellRanger-v7.1.0 and v9.0.1. Raw single-cell count matrices for all samples were imported using Seurat version 5 in a renv-managed R environment (R v4.5.0).

***Ambient RNA assessment and correction***

Ambient RNA was evaluated from paired raw and filtered count matrices using SoupX(v1.6.2). Empty droplets were defined as barcodes with <100 total UMIs (unique molecular identifiers), and background expression profiles were computed across these droplets. Contamination fractions were estimated using SoupX (SoupChannel, setClusters, autoEstCont) with quick clustering (scran::quickCluster) to define groups for estimation. For the iE7.5$^{Early}$-aE16.5 dataset, ambient RNA correction was applied, SoupX-corrected count matrices were generated by first building a Seurat object from CellRanger filtered counts, normalisation and scaling, PCA, a k-nearest neighbor (k-NN) graph, Louvain clustering at resolution 0.5, and using these clusters as SoupX groups; corrected counts were generated with adjustCounts and exported in 10x Genomics v3 format using DropletUtils::write10xCounts.

***Doublet detection and removal***

Doublets were detected per sample using DoubletFinder (v2.0.6). Low-quality cells expressing <2000 genes, <2000 UMIs, or > 10% mitochondrial genes were excluded before downstream analysis. Potential doublets expressing >8000 genes and >50000 gene counts were filtered out.

For each sample, preliminary preprocessing was performed (NormalizeData, variable feature selection with 3000 features, ScaleData, PCA, UMAP, a k-NN graph, Louvain clustering at resolution 0.8) to obtain group labels used for homotypic doublet estimation. DoubletFinder parameters were selected by sweeping pK (paramSweep/summarizeSweep/find.pK) and choosing the value maximizing the BC metric; the expected number of doublets was set from a sample-specific expected doublet rate and adjusted by the estimated homotypic proportion (modelHomotypic). DoubletFinder was run using PCs 1:30 with pN = 0.25, and cells classified as doublets were excluded from downstream analyses.

***Normalization, dimensionality reduction, integration, and clustering***

Normalization and variance stabilization was performed using SCTransform (v0.4.3), regressing out the mitochondrial percentage. The top 3000 HVGs excluding the sex-associated genes (*Xist*, *Gm13305*, *Tsix*, *Gm8730, Eif2s3y*, *Ddx3y*, *Uty, Kdm5d*) were used for PCA. Cells were clustered using a k-NN graph and Leiden clustering, followed by visualization using UMAP. Batch correction was performed using the first 30 PCs for Harmony Integration. The resulting Harmony reduction was used for a k-NN graph, followed by Leiden clustering and UMAP visualization.

*Post hoc* analysis was performed to remove residual poor-quality cells that were clustered by high mitochondrial percentages, as well as potential doublets identified by high total UMIs, high total gene number, and co-expression of markers genes from distinct lineages. The remaining cells were processed with an additional round of clustering for the final datasets of 5,356 cells for the iE7.5-aE16.5 dataset, and 21,749 cells for the iE7.5$^{Early}$-aE16.5 dataset. Each cell was assigned a cell-cycle state using the CellCycleScoring function. The ENS cell population from the iE7.5-aE16.5 dataset was subsequently subsetted and processed with an additional round of normalisation and clustering.

##### *Cell-type annotation*

Cell clusters were manually annotated based on assessing differentially expressed genes identified from the Wilcoxon rank-sum test (FindAllMarkers) implemented in Seurat, together with the expression of canonical markers genes for gut cell types. Cell type annotations were also benchmarked against predictions generated by an automated annotation tool - Single Cell Signature Scorer for Annotation (SCSA) (5) with hyperparameters LFC = 1.5 and p-value = 0.05.

##### *CloneID extraction*

We extracted the cell IDs from all cells of the same embryo that passed the quality control, and exported the cell IDs as a TSV file for CloneID extraction implemented in TREX pipeline (4). TREX uses reads from filtered cells, based on imported cell IDs, to align to the tdTomato-N transgene, and barcodes were recovered from alignment spanning the N region. Overrepresented barcodes identified from lentiviral barcode library sequencing and characterization (3) were excluded from CloneID construction. Barcodes retrieved with fewer than 7 characters were also removed. Reads with identical barcode, UMI and cell ID are collapsed into a consensus sequence. To error-correct barcode sequences, they are single-linkage clustered using a Hamming distance of 5. Cells were assigned the same CloneID when they shared both the same barcode number and identical barcode sequences (Jaccard index of 1.0, ie we applied the most stringent criteria for clone assignment).

##### *Clonal composition and clonal coupling analysis*

CloneIDs were mapped onto the Seurat metadata of each dataset using matched cell IDs to annotate cells as 'with CloneID' or 'without CloneID', and where applicable, assigned a corresponding CloneID (clone_nr). Clone size was defined as the number of cells sharing the same CloneID. Clones were categorized as singleton (one cell), single-cell-type multicellular (≥2 cells belonging to one annotated cell type), or multi-cell-type multicellular (≥2 cells belonging to more than one annotated cell type). For clonal composition analyses, cells were annotated to fibroblasts, pericytes, ICC, smooth muscle cells, myofibroblasts, mesothelial cells, endothelial cells, simple epithelium, foregut basal epithelium, ENS, and immune cells. Clone sharing across annotated cell types and gut regions was summarized using UpSet plots and chord diagrams. Regionally dispersed clones were determined by clones with cells present across distinct regions.

To quantify over- or under-representation of cell types sharing clones, clonal coupling scores were calculated as previously described (6, 7). Observed clone sharing was compared to randomised data, yielding a z-score between each pair of annotated cell types. The resulting clonal coupling z-scores represent how many standard deviations the observed clone sharing between each cell type pair is away from the mean of randomised data. Positive values indicate higher than expected clone sharing and negative values indicate lower than expected clone sharing. Analyses were performed using 1,000 random shuffles for multiple times.

To identify annotated cell types with similar clonal coupling patterns, Pearson correlations of z-scores were computed by which cell types are grouped by similarities in clonal coupling z-score profiles in complete lineage hierarchical clustering using Euclidean distance. Positive correlation values indicate that two cell types exhibit similar patterns of clonal coupling with other cell types, whereas negative correlation values indicate dissimilar clonal coupling patterns.

**REFERENCES (supplementary material and methods):**

### Supplementary Figure 1

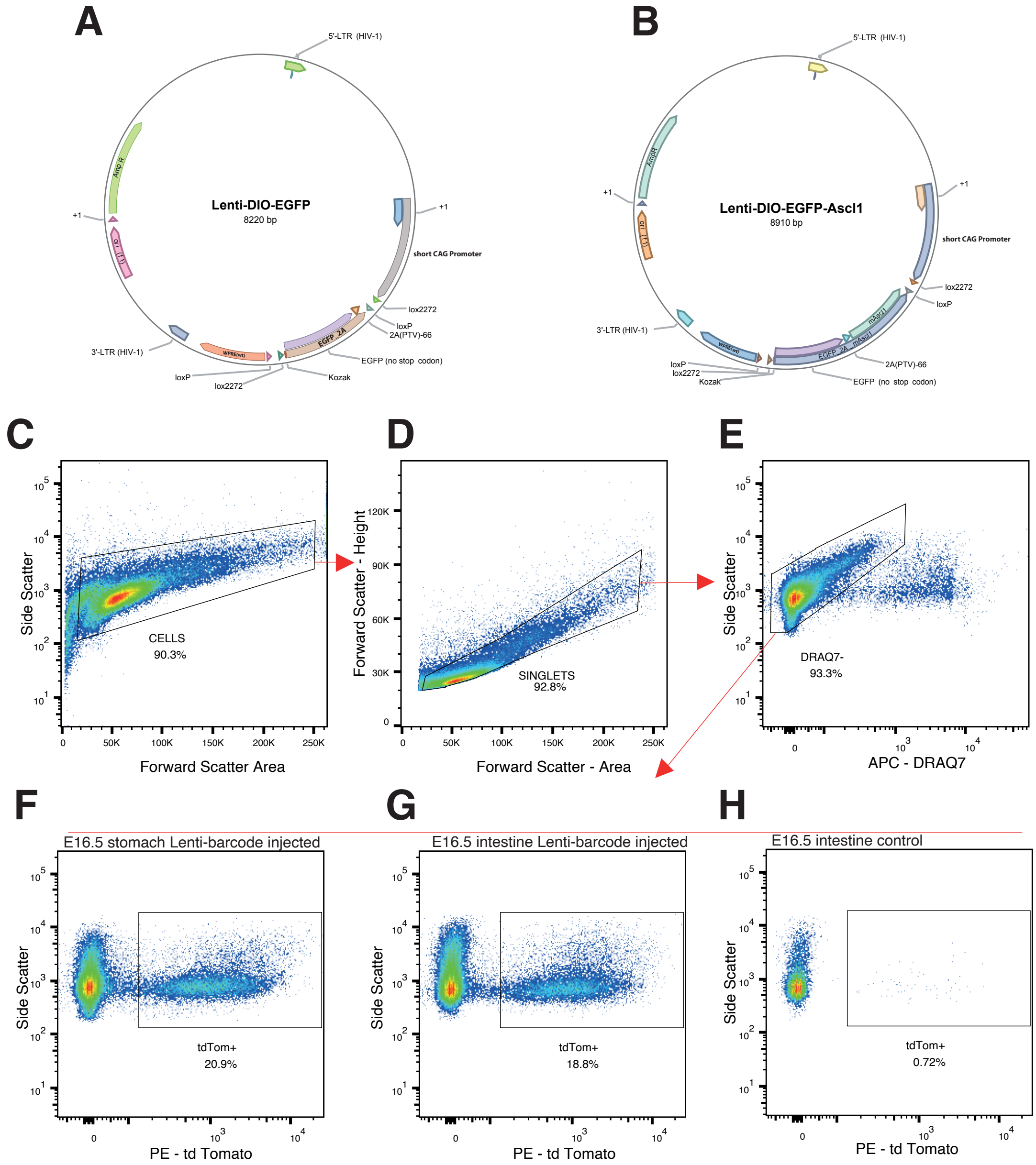


**Figure S1. Lentiviral vector design and flow cytometry gating strategy.**
**(A)** Plasmid map of Lenti-DIO-EGFP, a third-generation HIV-1–based lentiviral vector driving Cre-dependent EGFP expression via a shortened CAG promoter. The bicistronic cassette contains EGFP, and a PTV-1 2A self-cleaving peptide, flanked by double-floxed inverted orientation (DIO) for Cre-dependent expression. Transgene stability is conferred by WPRE, with vector maintenance supported by an f1 ori, AmpR selection cassette, and polyadenylation signal within the backbone.**(B)** Plasmid map of LV-DIO-EGFP-Ascl1, which shares the same backbone as **(A)** but encodes a EGFP–P2A–mAscl1 cassette, enabling Cre-dependent co-expression of EGFP and the transcription factor mAscl1 as separate proteins via PTV-1 2A-mediated self-cleavage. **(C-H)** Representative Flow Cytometry Data plots showing gating strategy for sorting tdTomato+ cells from E16.5 mouse gut injected with LV-EF1A-H2B-tdTomato-30N. Forward vs Side Scatter plot gating to identify cells **(C)**, Side Scatter vs Trigger Pulse Width to exclude doublets and multiplets **(D)**. DRAQ7- gating to exclude unhealthy cells **(E)**. Red fluorescence (580/30) used to distinguish tdTomato+ cells from autofluorescent cells in stomach or intestine of an injected embryo **(F, G)**. Gating of a control intestine **(H)**.

### Supplementary Figure 2

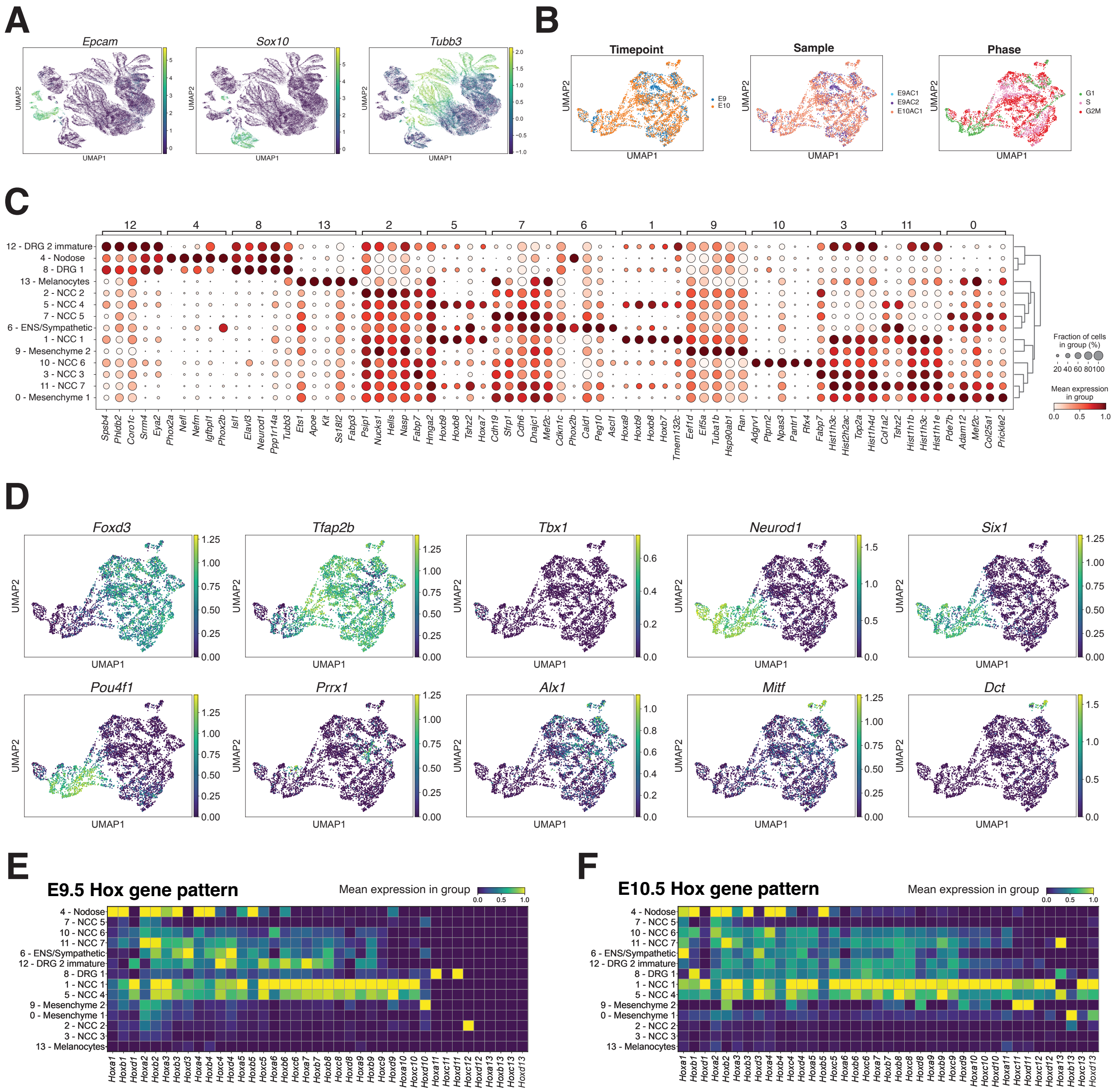


**Figure S2. Supportive data related to Figure 1 on neural crest/placode derived cells at E9.5-10.5 labelled by lentiviral nano-injection at E7.5 (A)** Feature plots of representative marker genes for epithelial cells (*Epcam*+), neural crest (*Sox10*+) and neurons (*Tubb3*+). **(B)** UMAP plot of annotated neural crest/placode-derived cells, colored by developmental timepoint, sample identity, and cell-cycle phase. **(C)** Dot plot displaying expression of the top five differentially expressed genes for each cluster. Dot size represents the fraction of cells expressing the gene, and color indicates normalized mean expression. **(D)** Feature plots of selected marker genes for neural crest (*Foxd3*+, *Tfap2b*+) and distinct neural crest/placode lineages, including nodose, DRG, cranial mesenchymal, and melanocyte populations. Heatmaps showing cluster expression of Hox genes at E9.5 **(E)** and E10.5 **(F).**

### Supplementary Figure 3

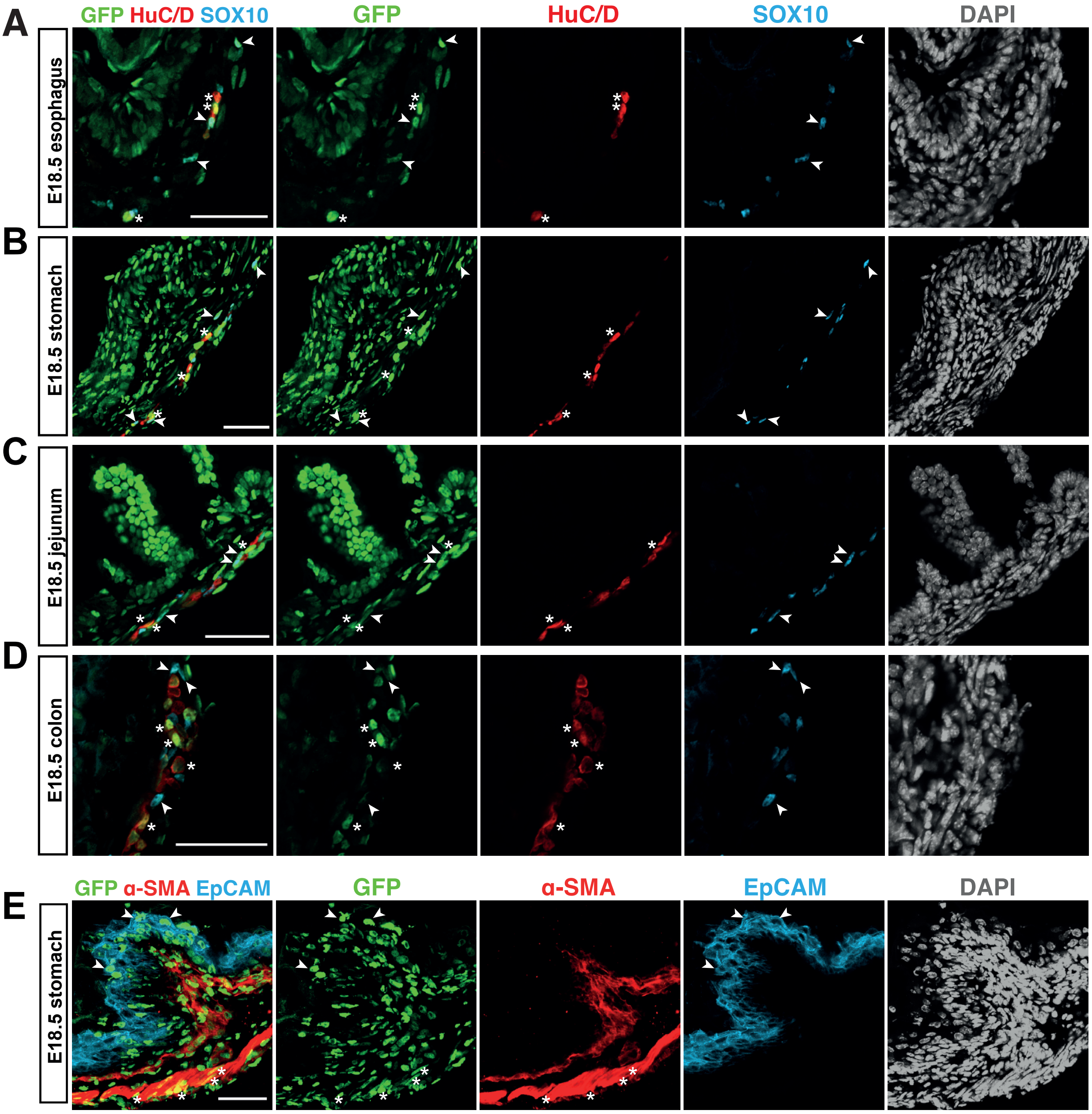


**Figure S3.** ***In utero*** **transduction of LV-H2B-GFP at E7.5 result in widespread GFP expression of the ENS and gut tube across regions. (A-D)** Representiative immunofluorescent images showing GFP expression in HuC/D+ neurons (asteriks) and SOX10+ progenitors/glia (arrowheads) in different gut regions. **(E)** Representiative immunofluorescent image of stomach showing GFP expression in α-SMA+ smooth muscle cells (asterisks) and EpCAM+ epithelial cells (arrowheads).Scale bars: 50$\mu$m.

### Supplementary Figure 4

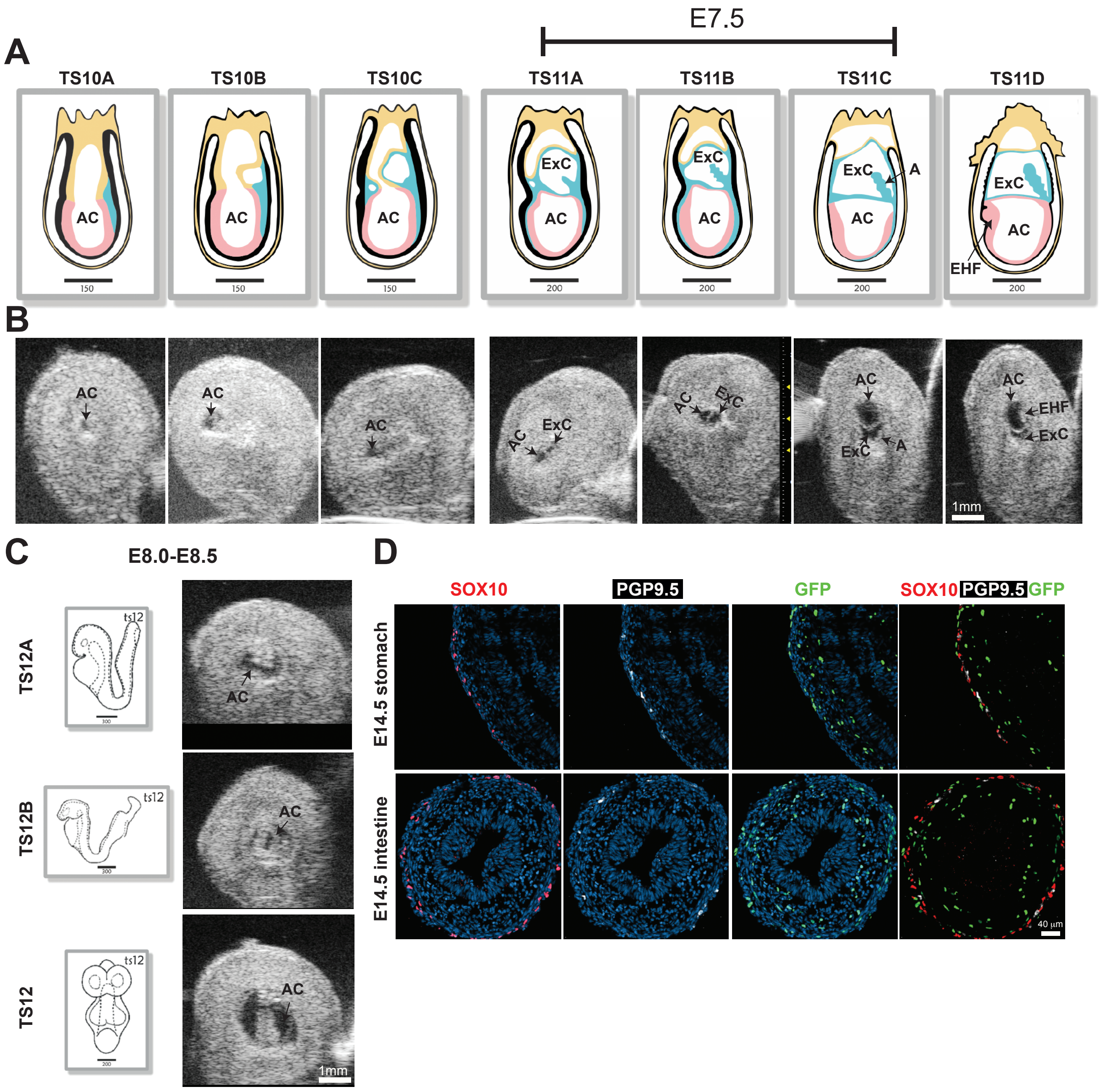


**Figure S4. Schematic drawings and representative ultrasound images of mouse Theiler stages.**
**(A)** Schematic drawings of E7.0-E7.75 mouse embryos showing different embryonic structures. TS11a-11c represent approximately the E7.5 stage. Images adapted from the eMouseAtlas Project (www.emouseatlas.org). **(B)** Corresponding ultrasound images of E7.0-E7.75 mouse embryos with structures labelled. Scale bar: 1mm. **(C)** Schematic drawings from the eMouseAtlas Project of TS12A-C, representing approximately E8.0-E8.25. Representative ultrasound images of E8.0-E8.5 mouse embryos. Scale bar: 1mm. **(D)** Representative confocal images of E14.5 stomach and intestine transduced with LV-H2B-GFP at E8.0-8.5 showing only sparse labelling of mesenchyme, and no epithelia nor ENS. Scale bar: 40$\mu$m.
AC: amniotic cavity; ExC: extracoelomic cavity; A: allantois and EHF: early head fold

### Supplementary Figure 5

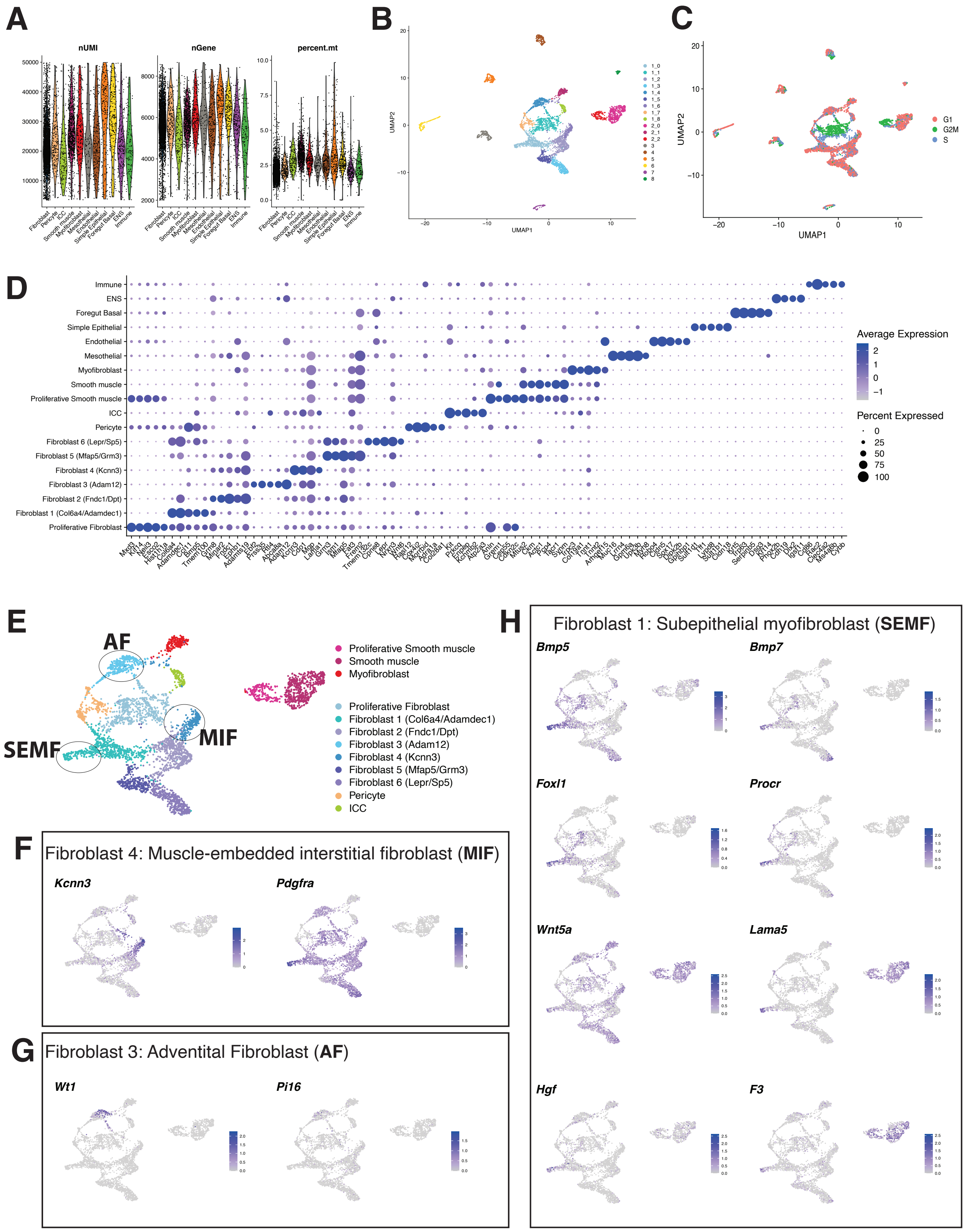

**Figure S5. Quality control and marker gene analysis of gut cell types in the iE7.5-aE16.5 dataset. (A)** Violin plots showing per-cell quality control metrics for each major cell type after filtering, including total reads (nUMI), number of unique genes detected (nGene), and percentage of mitochondrial transcripts (percent.mt). **(B)** UMAP plot of cells colored by unsupervised cluster assignment. **(C)** UMAP plot of cells colored by cell cycle phase. **(D)** Dot plot displaying expression of the top five differentially expressed genes for each refined cell type. Dot size represents the fraction of cells expressing the gene, and color indicates relative mean expression. **(E)** UMAP plot showing subclusters within fibroblast-like and smooth muscle-like clusters. **(F)** Feature plots showing markers of Muscle-embedded Interstitial Fibroblasts (MIF) in Fibroblast 4. **(G)** Feature plots showing expression of genes associated with Adventitial Fibroblasts (AF) in Fibroblast 3. **(H)** Feature plots showing expression of genes associated with Subepithelial Myofibroblasts (SEMF) in Fibroblast 1.

### Supplementary Figure 6

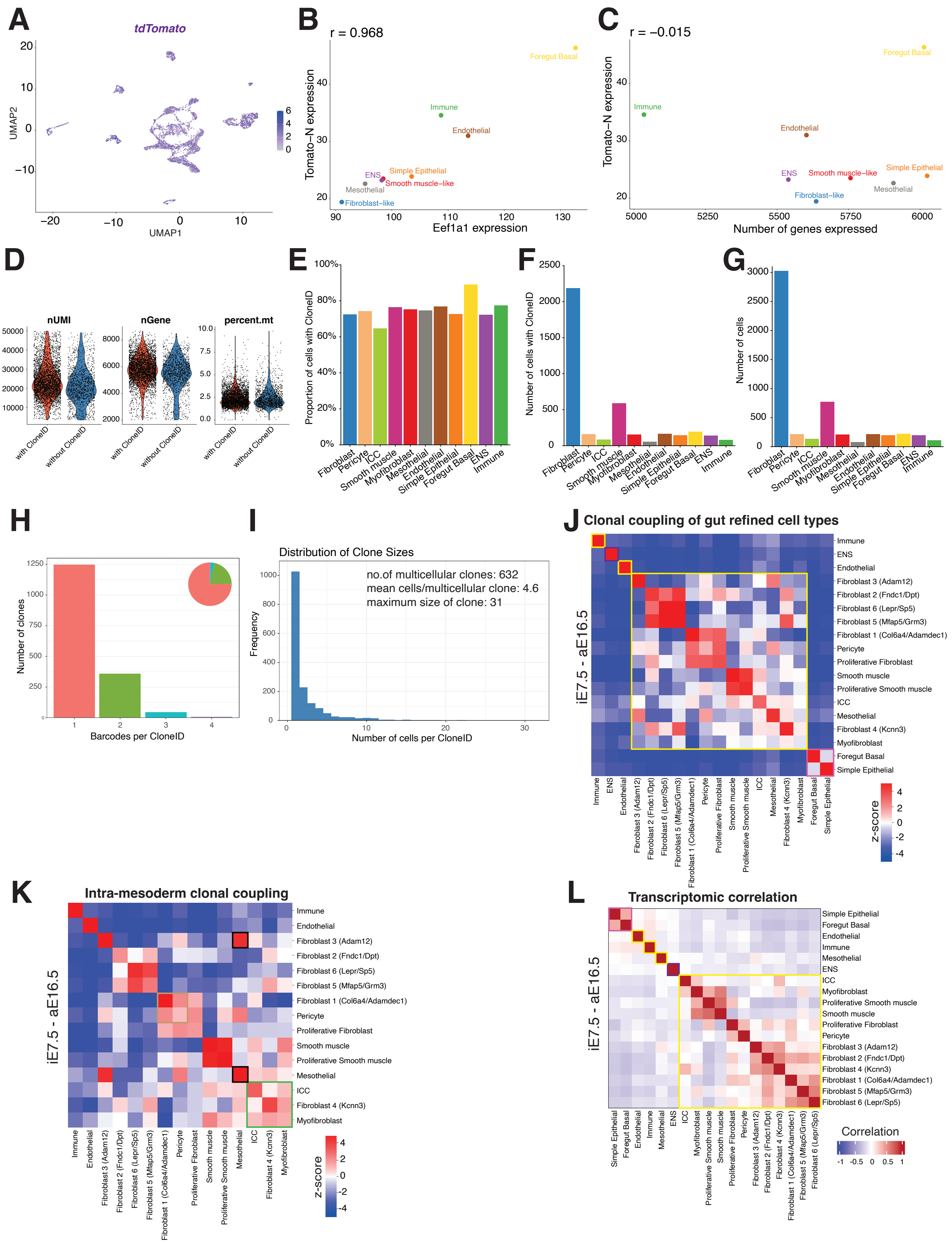

**Figure S6. CloneID assessment and clonal coupling analysis in the iE7.5-aE16.5 dataset.** **(A)** Feature plot displaying tdTomato expression. **(B)** Scatter plot showing a high Pearson correlation between Eef1a1 expression and tdTomato expression across major gut cell types, indicating that the synthetic Eef1a1 promoter recapitulates endogenous Eef1a1 expression patterns. **(C)** Scatter plot showing a low Pearson correlation between tdTomato expression and average number of genes detected per cell across major gut cell type, excluding library complexity as a confounder. **(D)** Violin plots showing per-cell quality control metrics for cells with and without CloneIDs. **(E)** Bar graph displaying the proportions of cells with CloneIDs per cell type. **(F)** Bar graph showing the number of cells with CloneIDs per cell type. **(G)** Bar graph showing the total number of cells per cell type. (H) Bar graph and pie chart illustrating the distribution of the number of unique barcodes per CloneID. **(I)** Bar graph showing distribution of the number of cells per CloneID, with summary statistics. **(J, K)** Heatmaps of clonal coupling z-scores, defined as the deviation in the number of shared CloneIDs between pairs of cell types in all clones (J) and clones with only mesoderm-derived cells (K) relative to randomized data, related to Figure 3F and G. **(L)** Heatmap showing transcriptomic correlations between refined cell types based on expression profiles. Colored boxes denote germ-layer origin of the cell type (J, L; pink: endoderm; purple: ectoderm; yellow: mesoderm) or lineage-coupled mesenchymal populations (K; green: Fibroblast 4 with myofibroblasts, ICC and smooth muscle cells; black: Fibroblast 3 with mesothelial cells; brown: Fibroblast 1 with pericytes).

### Supplementary Figure 7

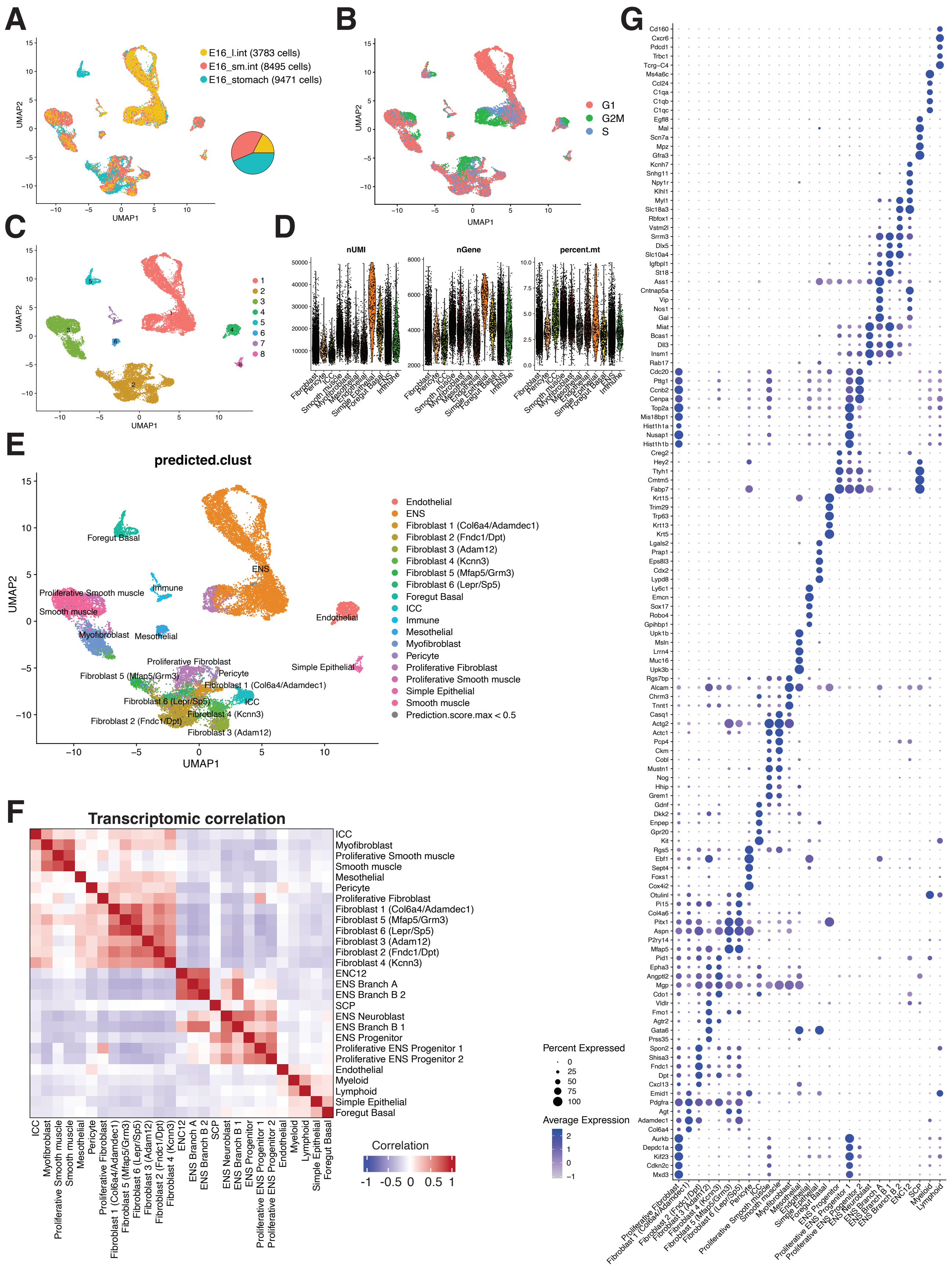

**Figure S7. Quality control and marker gene analysis of gut cell types in the iE7.5$^{Early}$-aE16.5 dataset. (A)** UMAP plot colored by sample origin. Pie chart indicates the proportions of cells contributed by each gut region (stomach, small intestine, large intestine). **(B)** UMAP plot colored by cell cycle phase. **(C)** UMAP plot colored by unsupervised cluster assignment. **(D)** Violin plots showing per-cell quality control metrics for each major cell type after filtering, including total reads (nUMI), number of unique genes detected (nGene), and percentage of mitochondrial transcripts (percent.mt). **(E)** UMAP plot showing predicted cell-type annotations by label transfer using the iE7.5-aE16.5 dataset as reference. **(F)** Heatmap showing transcriptomic correlations between refined cell types based on expression profiles. **(G)** Dot plot showing expression of top five differentially expressed genes across refined cell types. Dot size represents the percentage of cells expressing the gene, and color indicates relative mean expression.

### Supplementary Figure 8

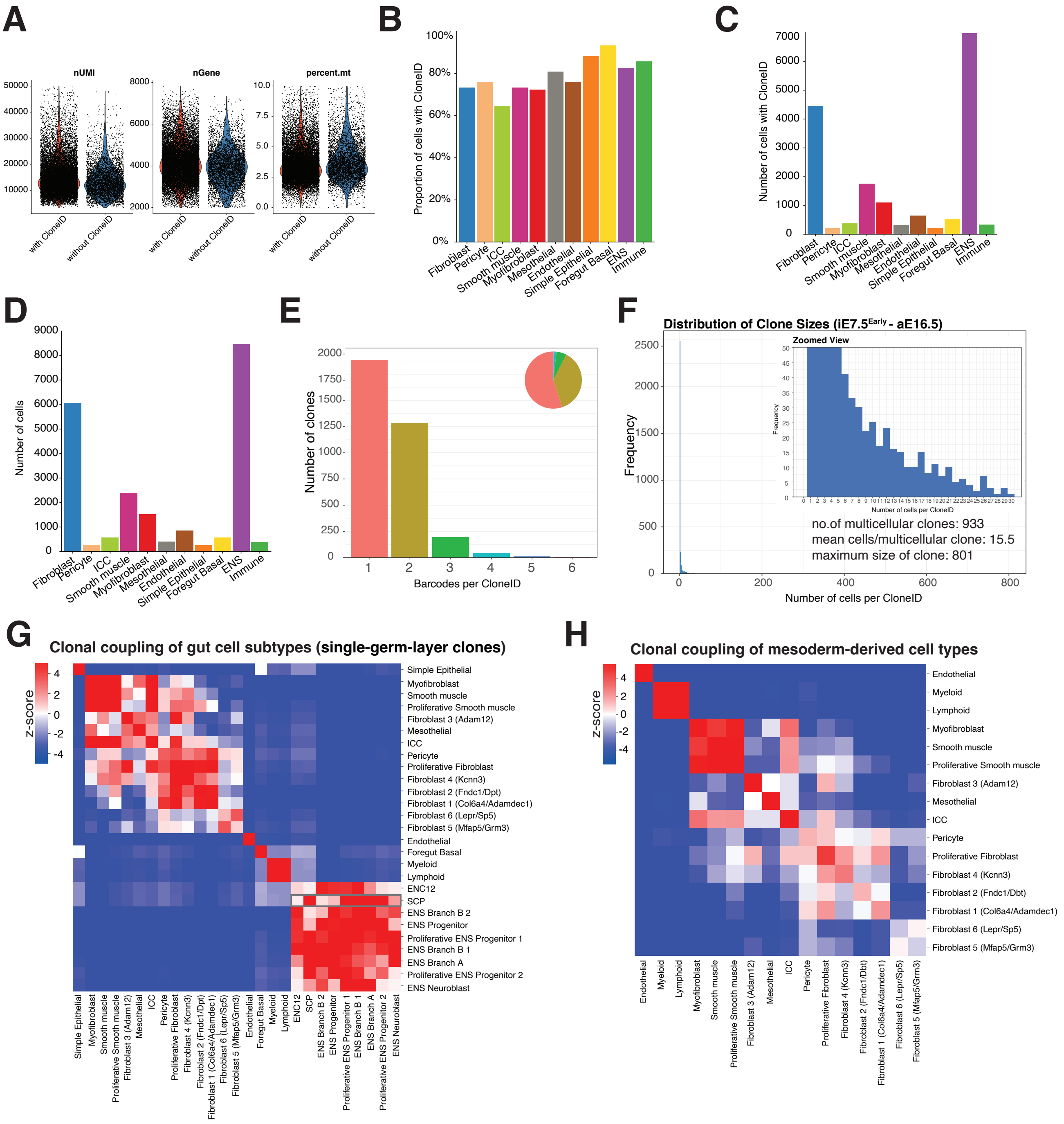

**Figure S8. CloneID assessment and clonal coupling analysis in the iE7.5$^{Early}$-aE16.5 dataset. (A)** Violin plots showing per-cell quality control metrics for cells with and without CloneIDs. **(B)** Bar graph displaying the proportions of cells with CloneIDs per cell type. **(C)** Bar graph showing the number of cells with CloneIDs per cell type. **(D)** Bar graph showing the number of cells per cell type. **(E)** Bar graph and pie chart illustrating the distribution of the number of unique barcodes per CloneID. **(F)** Bar graph showing the distribution of the number of cells per CloneID, with summary statistics. **(G,H)** Heatmaps of clonal coupling z-scores, defined as the deviation in the number of shared CloneIDs between pairs of cell types found in all single-germ-layer clones (G) and in single-germ-layer clones with only mesoderm-derived cells (H) relative to randomized data, related to Figure 4H, I. Gray box in (G) highlights clonal coupling between SCP and all ENS states, but relatively lower coupling to the two most mature neuron clusters (ENC12 and Branch B2) in agreement with the later arrival of SCP to the gut than of vagal neural crest. Note that Branch B1 and B2 are not equivalents of Branch B1 and B2 defined in Morarach et al, 2021.

### Supplementary Figure 9

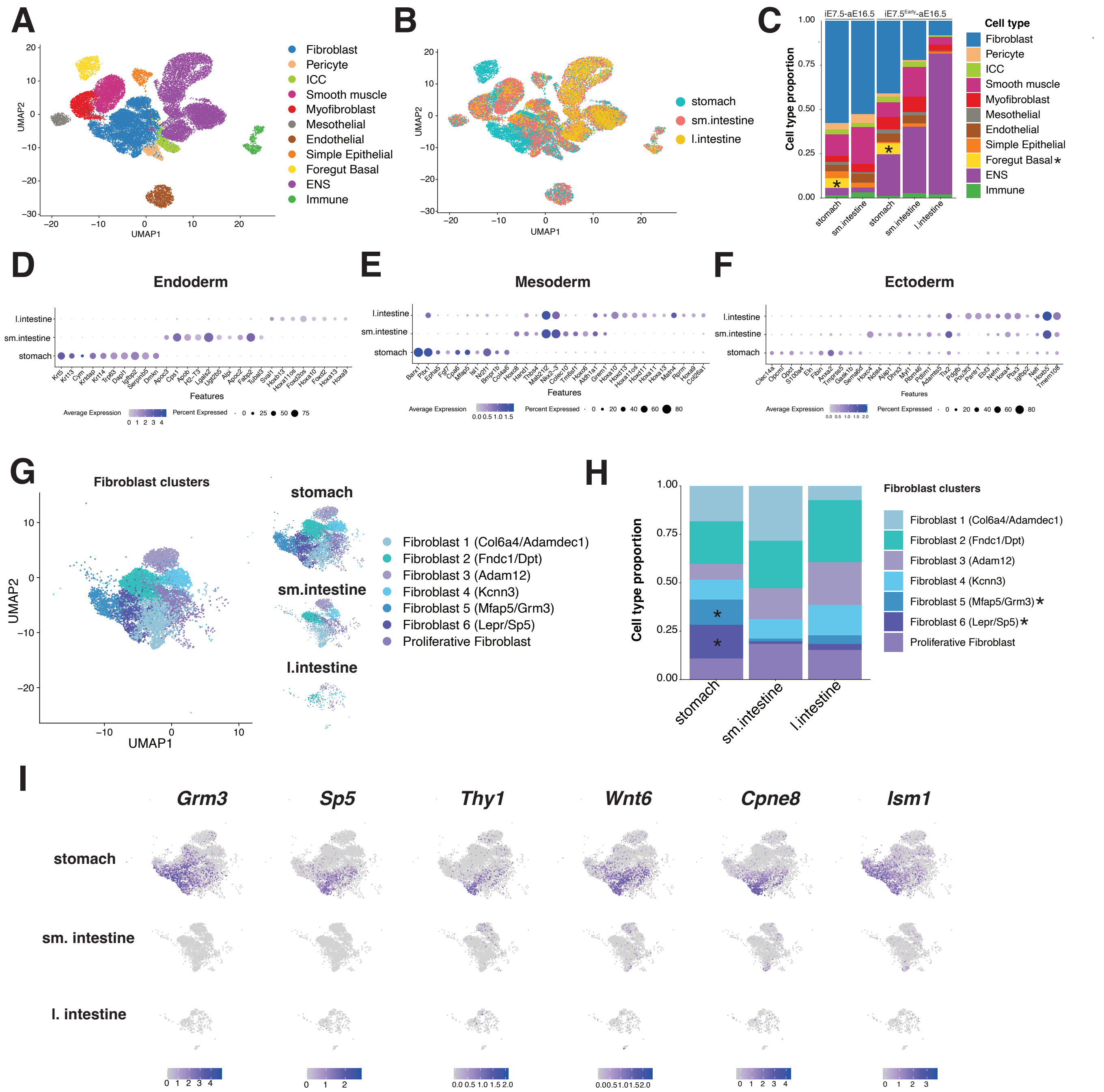


**Figure S9. Regional distribution of gut cell types and gene expression patterns. (A)** UMAP plot of integrated iE7.5-aE16.5 and iE7.5[Early]-aE16.5, showing major cell types. **(B)** UMAP plot colored by tissue of origin. **(C)** Stacked bar plots showing the proportion of major cell types across gut regions and samples. Asterisks indicate Foregut Basal cells, which are detected exclusively in stomach samples. **(D–F)** Dot plots showing the top 10 differentially expressed genes of cells derived from each germ layer: endoderm **(D)**, mesoderm **(E)**, and ectoderm **(F)**, across gut regions. Dot size represents the percentage of cells expressing each gene, and color intensity indicates average expression level. **(G)** UMAP plot showing fibroblast subclusters across gut regions (stomach, small intestine, and large intestine). **(H)** Stacked bar plots showing the proportion of fibroblast subclusters across gut regions. Asterisks indicate Fibroblast 5/6 enrichment in the stomach. **(I)** Feature plots showing regional expression patterns of Fibroblast 5/6 enriched genes across gut regions.

### Supplementary Figure 10

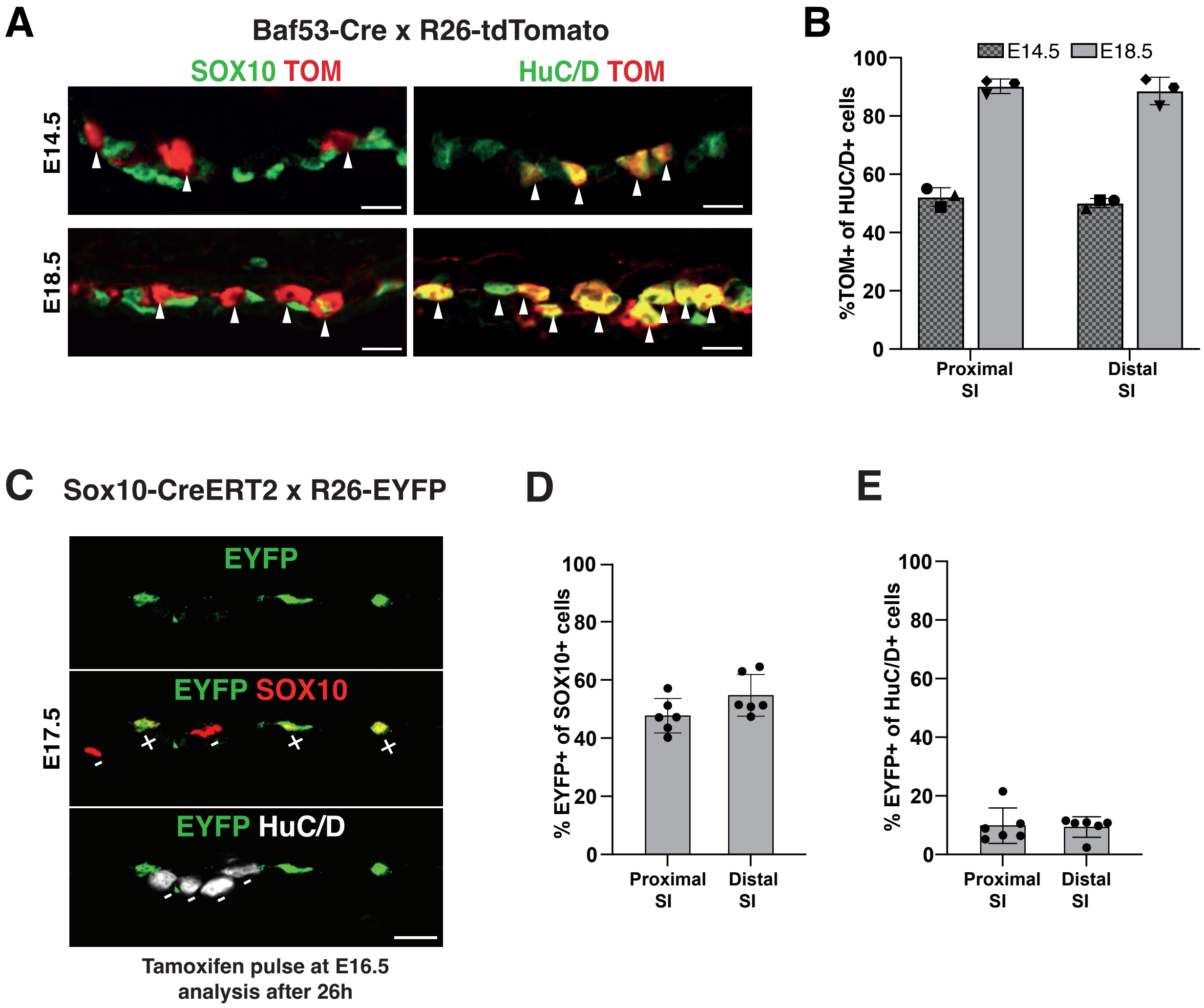


**Figure S10. Baf53b-Cre and Sox10-CreERT mice can be used to induce reporter expression in developing ENS neurons and progenitors/glia, respectively. (A)** Representative confocal images of small intestine showing TOM expression in HuC/D+ neurons but not in SOX10+ glia/progenitors. TOM+ cells is indicated with white arrowheads. Scale bars: 20$\mu$m. **(B)** Bar graphs showing percentage of HuC/D+ cells expressing TOM as represented in (A). Bars represent mean ± SD. n=3 mice **(C)** Representative confocal images of small intestine showing that EYFP is predominantly induced in SOX10+ cells rather than HuC/D+ cells after 26 hours following tamoxifen administration in Sox10-CreERT2; R26-EYFP females at E16.5. (D) Bar graph showing percentage of SOX10 cells expressing EYFP as represented in (C). **(D)** Bar graph showing percentage of HuC/D cells expressing EYFP as represented in (C). Bars represent mean ± SD. n=6 mice. SI: small intestine.
